# Benchmarking Single-Cell RNA Sequencing Protocols for Cell Atlas Projects

**DOI:** 10.1101/630087

**Authors:** Elisabetta Mereu, Atefeh Lafzi, Catia Moutinho, Christoph Ziegenhain, Davis J. MacCarthy, Adrian Alvarez, Eduard Batlle, Sagar, Dominic Grün, Julia K. Lau, Stéphane C. Boutet, Chad Sanada, Aik Ooi, Robert C. Jones, Kelly Kaihara, Chris Brampton, Yasha Talaga, Yohei Sasagawa, Kaori Tanaka, Tetsutaro Hayashi, Itoshi Nikaido, Cornelius Fischer, Sascha Sauer, Timo Trefzer, Christian Conrad, Xian Adiconis, Lan T. Nguyen, Aviv Regev, Joshua Z. Levin, Swati Parekh, Aleksandar Janjic, Lucas E. Wange, Johannes W. Bagnoli, Wolfgang Enard, Marta Gut, Rickard Sandberg, Ivo Gut, Oliver Stegle, Holger Heyn

## Abstract

Single-cell RNA sequencing (scRNA-seq) is the leading technique for charting the molecular properties of individual cells. The latest methods are scalable to thousands of cells, enabling in-depth characterization of sample composition without prior knowledge. However, there are important differences between scRNA-seq techniques, and it remains unclear which are the most suitable protocols for drawing cell atlases of tissues, organs and organisms. We have generated benchmark datasets to systematically evaluate techniques in terms of their power to comprehensively describe cell types and states. We performed a multi-center study comparing 13 commonly used single-cell and single-nucleus RNA-seq protocols using a highly heterogeneous reference sample resource. Comparative and integrative analysis at cell type and state level revealed marked differences in protocol performance, highlighting a series of key features for cell atlas projects. These should be considered when defining guidelines and standards for international consortia, such as the Human Cell Atlas project.

---

Single-cell genomics provides an unprecedented view of the cellular makeup of complex and dynamic systems. Single-cell transcriptomics approaches in particular have led the technological advances that allow unbiased charting of cell phenotypes^1^. The latest improvements in single-cell RNA sequencing (scRNA-seq) allow these technologies to scale to thousands of cells per experiment, providing comprehensive profiling of cellular composition^2,3^. This has led to the identification of novel cell types and the fine-grained description of cell plasticity in dynamic systems, such as development^4,5^. The latest large-scale efforts are attempting to produce cellular maps of entire cell lineages, organs and organism^6,7^, with probably the most notable effort being the initiation of the Human Cell Atlas (HCA) project^8^. To comprehensively chart the cellular composition of the human body, the HCA project conducts phenotyping at the single-cell level. It will advance our understanding of tissue function and serve as a reference to pinpoint variation in healthy and disease contexts. In addition to methods that capture the spatial organization of tissues^9,10^, the main approach to create a first draft human cell atlas is scRNA-seq-based transcriptome analysis of dissociated cells, in which tissues are disaggregated and individual cells are captured by cell sorting or using microfluidic systems^1^. In sequential processing steps, the RNA is reverse transcribed to cDNA, amplified and processed to sequencing-ready libraries. Continuous technological development has improved the scale, accuracy and sensitivity of the initial scRNA-seq methods, and now allows us to create tailored experimental designs by selecting from a plethora of different scRNA-seq protocols. However, there are marked differences between these methods, and it is still not clear which are the best protocols for drawing a cell atlas.

Experience from other large-scale consortium efforts has shown that neglecting benchmarking, standardization and quality control at the beginning can lead to major problems later on in the project, when investigators are attempting to exploit the results^11^. The overall success of any project depends critically on bringing the work of different consortium partners up to a high common standard. Thus, before launching into large-scale data collection efforts for the HCA and similar projects, it is important to conduct a comprehensive comparison of available single-cell profiling techniques.

In this paper, to extend current efforts to compare the molecule capture efficiency of scRNA-seq methods^12,13^, we have systematically evaluated the power of these techniques to describe tissue complexity, and their suitability for building a cell atlas. We performed a multi-center benchmarking study to compare the most common scRNA-seq protocols using a unified reference sample resource. By analyzing human peripheral blood and mouse colon tissue, we have covered a broad range of cell types and states, in order to represent common scenarios in cell atlas projects. We have also added spike-in cell lines to allow us to assess sample composition, and have combined different species to pool samples into a single reference. We performed a comprehensive comparative analysis of 13 different scRNA-seq protocols, representing the most commonly used methods. We applied a wide range of different quality control metrics to evaluate datasets from different perspectives, and to test their suitability for producing a reproducible, integrative and predictive reference cell atlas.

## Results

### Reference sample and experimental design

A variety of scRNA-seq methods have been developed, and their utility proven, in single-cell transcriptome analysis of complex and dynamic tissues. The available protocols vary in the efficiency of RNA molecule capture, resulting in differences in sequencing library complexity and sensitivity to identify transcripts and genes^12–14^. However, there has been no systematic testing of how their performance varies between cell types, and how this affects the resolution of cellular phenotyping of complex samples. To address this problem, we benchmarked current scRNA-seq protocols to inform the methodology selection process of cell atlas projects. Ideally, methods should a) be accurate and free of technical biases, b) be applicable across distinct cell properties, c) fully disclose tissue heterogeneity, including subtle differences in cell states, d) produce reproducible expression profiles, e) comprehensively detect population markers, f) be integrable with other methods, and g) have predictive value with cells mapping confidently to a reference atlas.

To perform a systematic comparison of scRNA-seq methods for cell atlas projects, we created a reference sample containing: i) a high degree of cell type heterogeneity with various frequencies, ii) closely related subpopulations with subtle differences in gene expression, iii) a defined cell composition with trackable markers, and iv) cells from different species. For this study, we selected human peripheral blood mononuclear cells (PBMC) and mouse colon, which are tissue types with highly heterogeneous cell populations, as determined by previous single-cell sequencing studies^15,16^. In addition to the well-defined cell types, both tissues contain cells in transition states that present subtle transcriptional differences. These tissues also have a wide range of cell sizes and RNA contents, which are key parameters that affect performance in cell capture and library preparation. Interrogating tissues from different species allowed us to pool samples and exclude cell doublets. In addition to the intra-sample complexity, the spiked-in cell lines enabled the identification of batch effects and biases introduced during cell capture and library preparation. We added cell lines with distinct fluorescent markers that allowed us to track them during sample preparation.

Specifically, the reference sample contained (% viable cells): PBMC (60%, human), colon (30%, mouse), HEK293T (6%, RFP labelled human cell line), NIH3T3 (3%, GFP labelled mouse cells) and MDCK (1%, TurboFP650 labelled dog cells) (**Figure 1**). To reduce variability due to technical effects during library preparation, the reference sample was prepared in a single batch, distributed into aliquots of 250,000 cells, and cryopreserved. We have previously shown that cryopreservation is suitable for single-cell transcriptomics studies of these tissue types^17^. For cell capture and library preparation, the thawed samples underwent FACS separation to remove damaged cells and physical doublets, except for the single-nucleus experiment.

**Figure 1.**
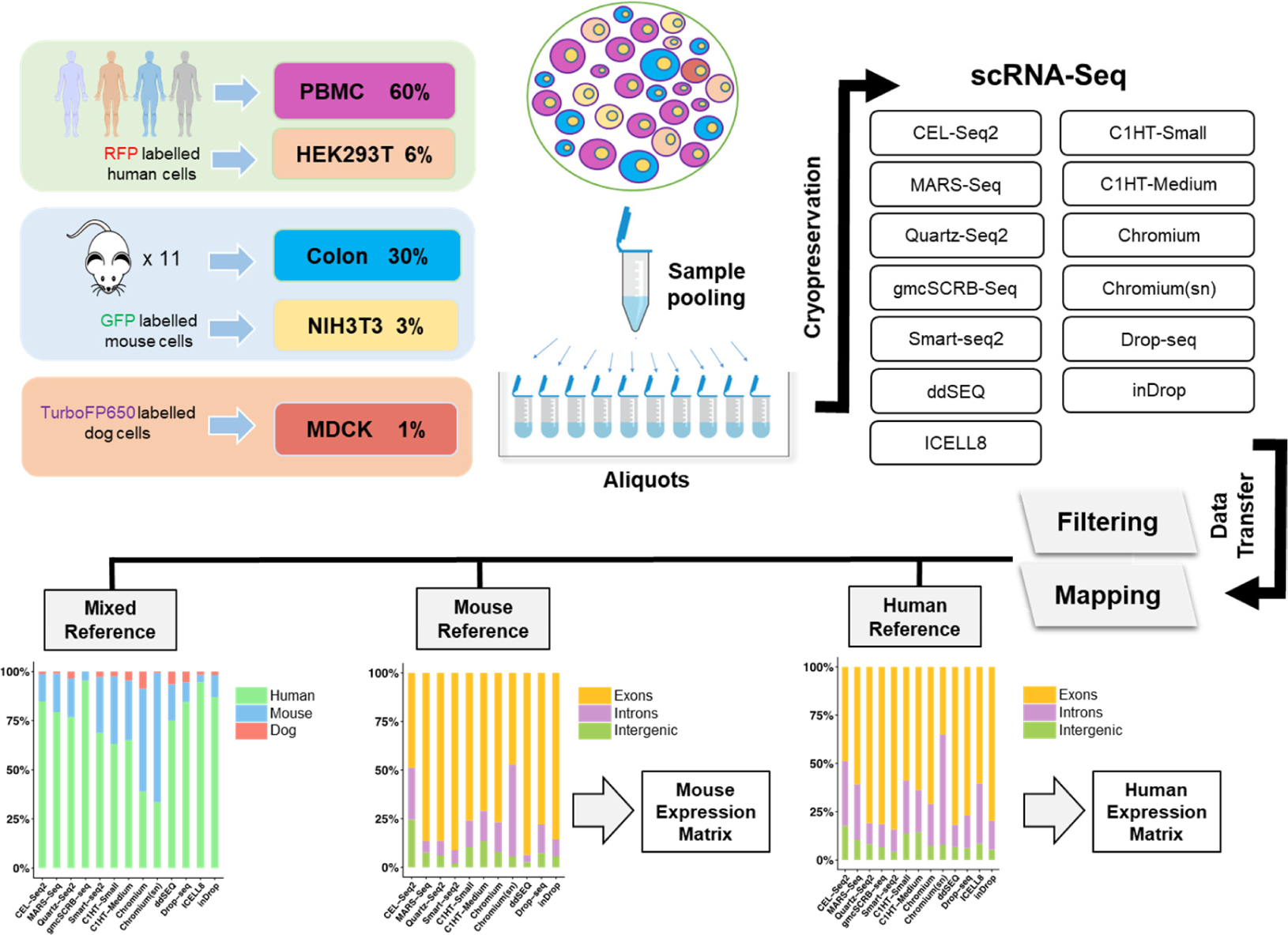
Overview of the experimental design and data processing. The reference sample consists of human PBMC (60%) and HEK293T (6%), mouse colon (30%) and NIH3T3 (3%) and dog MDCK (1%). The sample was prepared in one single batch, cryopreserved and sequenced by 13 different sc/snRNA-seq methods. Sequences were uniformly mapped to a joint human, mouse and canine reference and then separately to produce gene expression counts for each sequencing method.

### A reference dataset for benchmarking experimental and computational protocols

To obtain sufficient sensitivity to capture low-frequency cell types and subtle differences in cell state, we profiled ∼3,000 cells with each scRNA-seq method. In total, we produced datasets for 5 microtiter plate-based methods and 7 microfluidics systems, including cell-capture technologies based on droplets (4), nanowells (1) and integrated fluidic circuits (IFC), to capture small (1) and medium (1) sized cells (**Figure 1** and **Table S1**). We also included experiments to produce single-nucleus RNA sequencing (snRNA-seq) libraries (1), and an experimental variant that profiled >50,000 cells to produce a reference of our complex sample. The unified sample resource and standardized sample preparation (**Online Methods**) were designed to largely eliminate sampling effects, and allow the systematic comparison of scRNA-seq protocol performance.

To compare the different technologies, and to create a resource for the benchmarking and development of computational tools (e.g. batch effect correction, data integration and annotation), all datasets were processed in a uniform manner. Therefore, we designed a streamlined primary data processing pipeline tailored to the peculiarities of the reference sample (**Online Methods**). Briefly, raw sequencing reads were mapped to a joint human, mouse and canine reference genome and separately to their respective references to produce gene count matrices for subsequent analysis (data resource openly available). Consistent with the design of the reference sample, we detected most cells as human (63-95%) or mouse (4-34%; **Figure 1**). Notably, we observed a higher fraction of mouse colon cells in the single-nucleus sequencing dataset (Chromium (sn)). This could result from damaging the more fragile colon cells during sample preparation and resulting in proportionally fewer colon cells when selecting for cell viability. Indeed, when we skipped the viability selection step in the single-cell Chromium experiment as done in the single-nucleus experiment, we observed the same shift in composition towards mouse cells, suggesting that cell viability staining excludes cells that are amenable for scRNA-seq. Consequently, replacing viability staining with a thorough *in silico* quality filtering in cell atlas experiments might better conserve the composition of the original tissue. The canine cells, spiked-in at a low concentration, were detected by all protocols (1-9%) except gmcSCRB-seq. Furthermore, the different methods showed notable differences in mapping statistics between different genomic locations (**Figure 1**). As expected, due to the presence of unprocessed RNA in the nucleus, the snRNA-seq experiment detected the highest proportion of introns, although several scRNA-seq protocols also showed high frequencies of intronic and intergenic mappings.

### Molecule capture efficiency and library complexity

We produced reference datasets by analyzing 30,807 human and 19,749 mouse cells (Chromium V2; **Figure 2a-c**). The higher cell number allowed us to annotate the major cell types in our reference sample, and to extract population-specific markers (**Table S2**). Noteworthy, the reference samples solely provided the basis to assign cell identities and gene sets and was not utilized to quantify the methods’ performance. This strategy ensured that the choice of technology to derive the reference was not influencing downstream analyses. Indeed, cell clustering and reference-based cell annotation showed high agreement (average 80%; **Online Methods**) and only cells with consistent annotations were used subsequently for comparative analysis at cell type level. Notably, the PBMCs (human) and colon cells (mouse) represented two largely different scenarios. While the differentiated PBMCs clearly separated into subpopulations (e.g. T/B-cells, monocytes, **Figure 2b** and **Supplementary Fig. 1a, 2a-d**), colon cells were ordered as a continuum of cell states that differentiate from intestinal stem cells into the main functional units of the colon (i.e. absorptive enterocytes and secretory cells, **Figure 2c** and **Supplementary Figs. 1b, 3a-d**). After identifying major subpopulations and their respective markers in our reference sample, we clustered the cells of each sc/snRNA-seq protocol and annotated cell types using *matchSCore2* (**Online Methods**). This algorithm allows a gene marker-based projection of single cells (cell-by-cell) onto a reference sample and, thus, the identification of cell types in our datasets (**Supplementary Fig. 4** and **5**).

**Figure 2.**
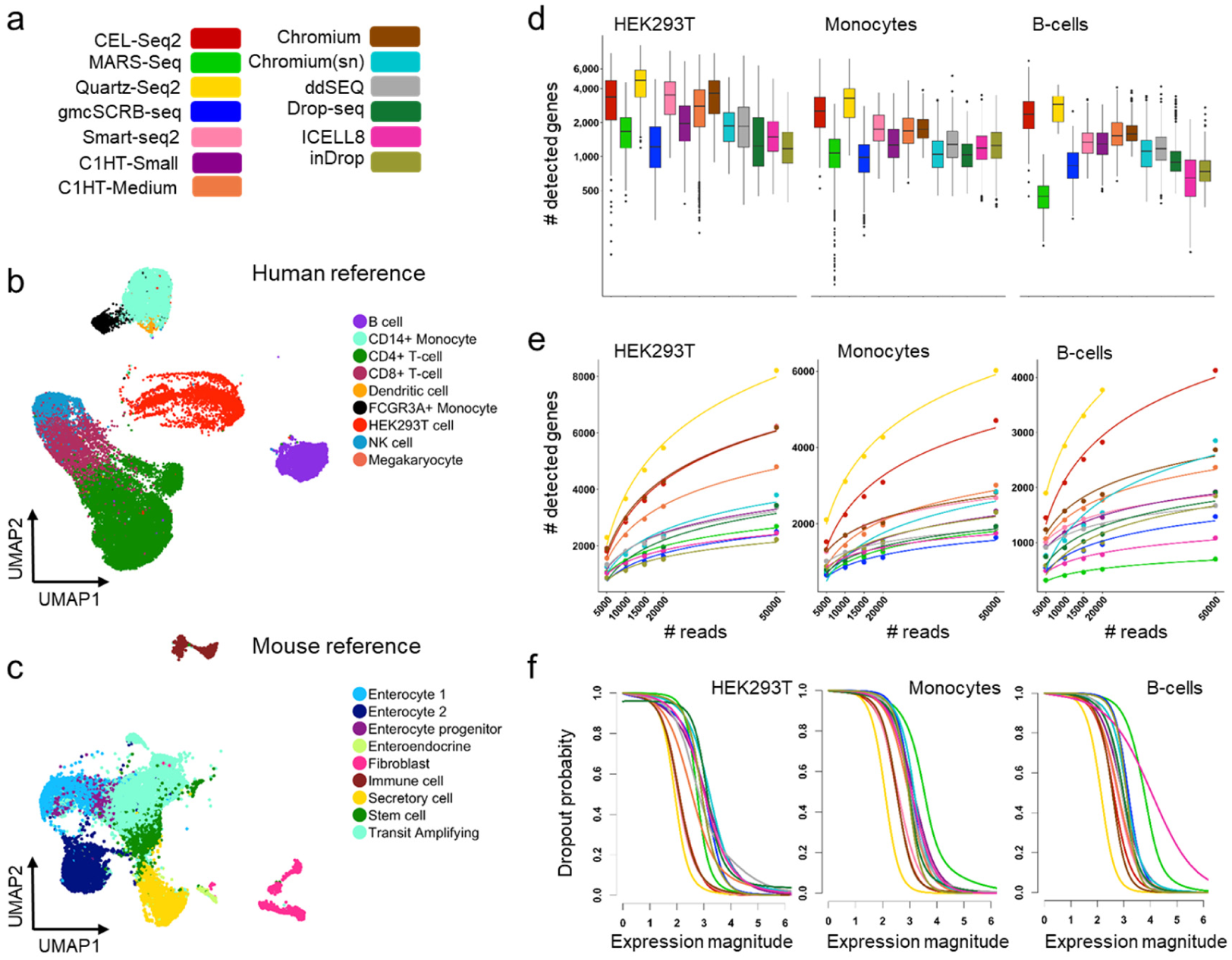
Comparison of 13 sc/snRNA-seq methods. **a.** Color legend of sc/snRNA-seq protocols. **b.** UMAP of 30,807 cells from the human reference sample (Chromium) colored by cell type annotation. **c.** UMAP of 19,749 cells from the mouse reference (Chromium) colored by cell type annotation. **d.** Boxplots comparing the number of genes detected across protocols, in downsampled (20K) HEK293T cells, monocytes and B-cells. Cell identities were defined by combining the clustering of each dataset and cell projection onto the reference. **e.** Number of detected genes at step-wise downsampled sequencing depths. Points represent the average number of detected genes as a fraction of all cells of the corresponding cell type at the corresponding sequencing depth. **f.** Dropout probabilities as a function of expression magnitude, for each protocol and cell type, calculated on downsampled data (20K).

**Figure 3.**
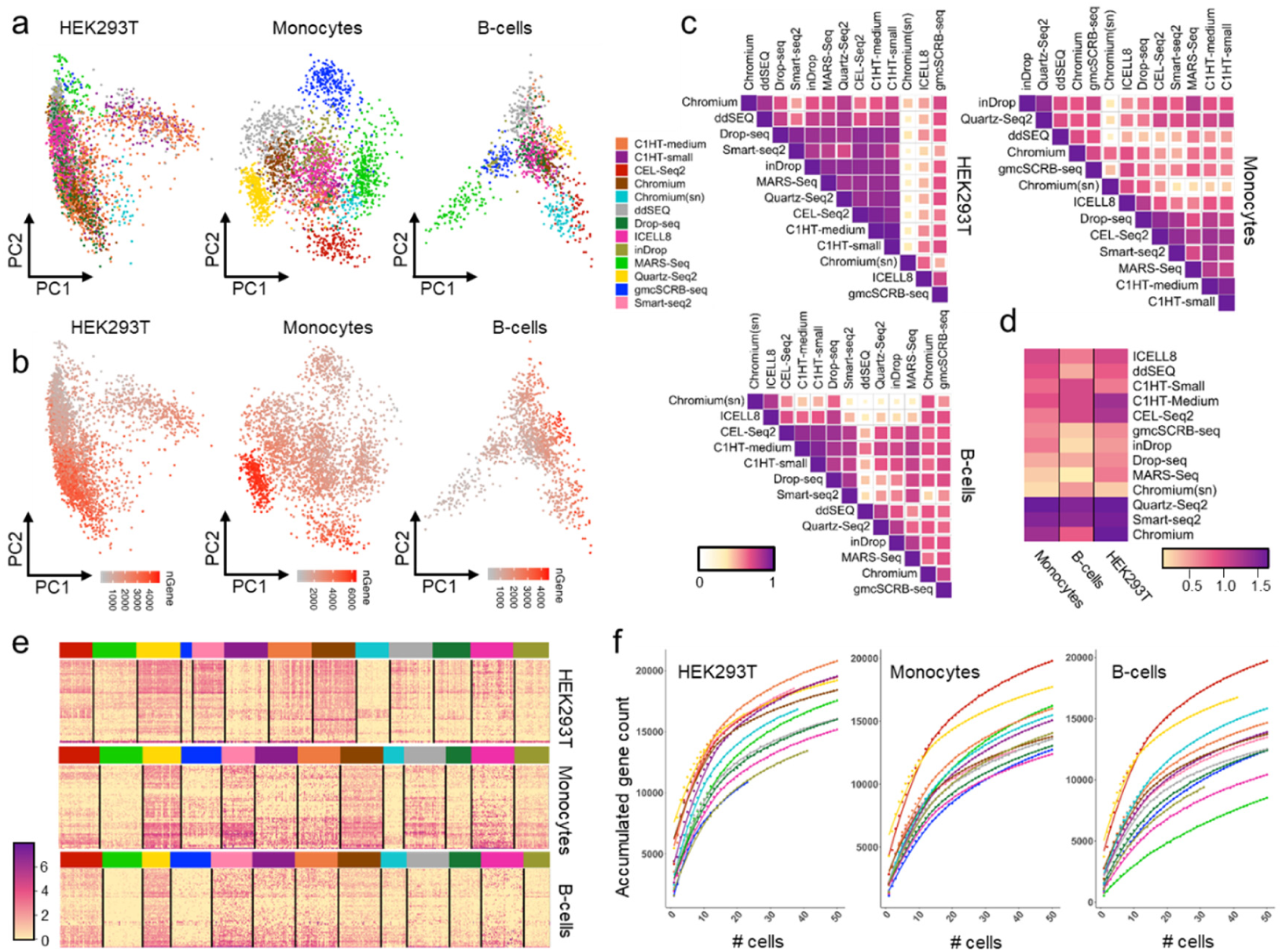
Similarity measures of sc/snRNA sequencing methods. **a,b.** PCA analysis on downsampled data (20K) using highly variable genes between protocols, separated into HEK293T cells, monocytes and B-cells, and color-coded by protocol (**a**) and number of detected genes per cell (**b**). **c.** Pearson correlation plots across protocols using expression of common genes. For a fair comparison, cells were downsampled to the same number for each method. Protocols are ordered by agglomerative hierarchical clustering. **d.** Heatmap representing average log expression values of downsampled (20K) HEK293T cell, monocyte, and B-cell reference markers per protocol. **e.** Heatmap representing the log expression values of HEK293T cell, monocyte and B-cell reference markers on downsampled data (20K). **f.** Cumulative gene counts per protocol as the average of 100 randomly sampled HEK293T cells, monocytes and B-cells separately on downsampled data (20K).

**Figure 4.**
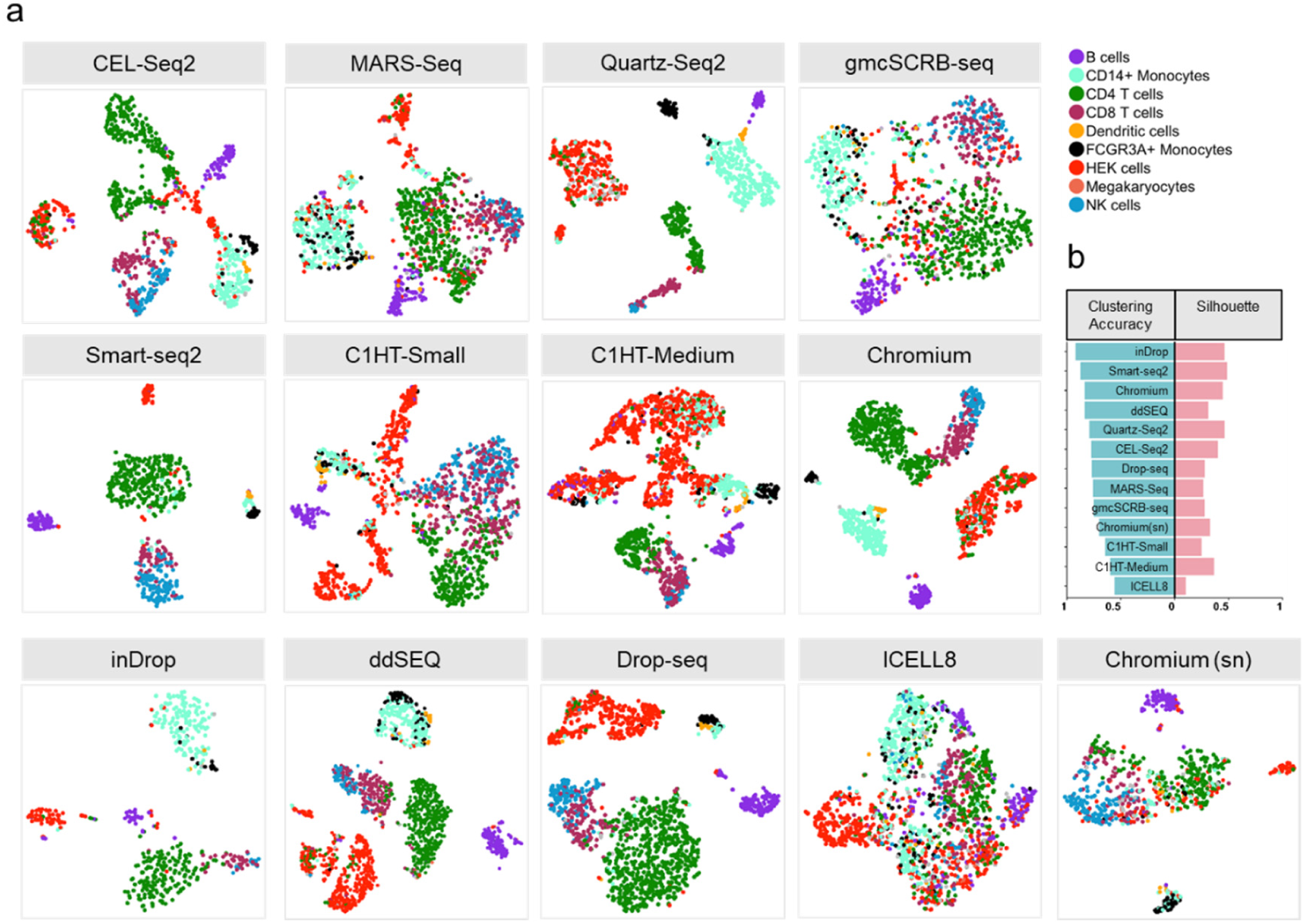
Clustering analysis of 13 sc/snRNA-seq methods on downsampled datasets (20K). **a.** T-SNE visualizations of unsupervised clustering in human samples from 13 different methods. Each dataset was analyzed separately after downsampling to 20K reads per cell. Cells are colored by cell type inferred by *matchSCore2* before downsampling. Cells that did not achieve a probability score of 0.5 for any cell type were considered unclassified. **b.** Clustering accuracy and Average Silhouette Width for clusters in each protocol.

To compare mRNA capture efficiencies among protocols we downsampled the sequencing reads per cell to a common depth and step-wise reduced fractions (100% to 25%). Library complexity was determined separately for largely homogenous cell types with markedly different cell properties and function, namely human HEK293T cells, monocytes and B-cells (**Figure 2d,e**), and mouse colon secretory and transit-amplifying (TA) cells (**Supplementary Fig. 6a,b**). We observed large differences in the number of detected genes between the protocols, with consistent trends across cell types and gene quantification strategies (**Supplementary Fig. 6c**). Notably, some protocols, such as Smart-seq2 and Chromium V2, performed better with higher RNA quantities (HEK293T) compared to lower starting amounts (monocytes and B-cells), suggesting an input-sensitive optimum. Consistent with the variable library complexity, the protocols presented large differences in drop-out probabilities (**Figure 2f**), with Quartz-seq2, Chromium V2 and CEL-seq2 showing consistently lower probability.

**Figure 5.**
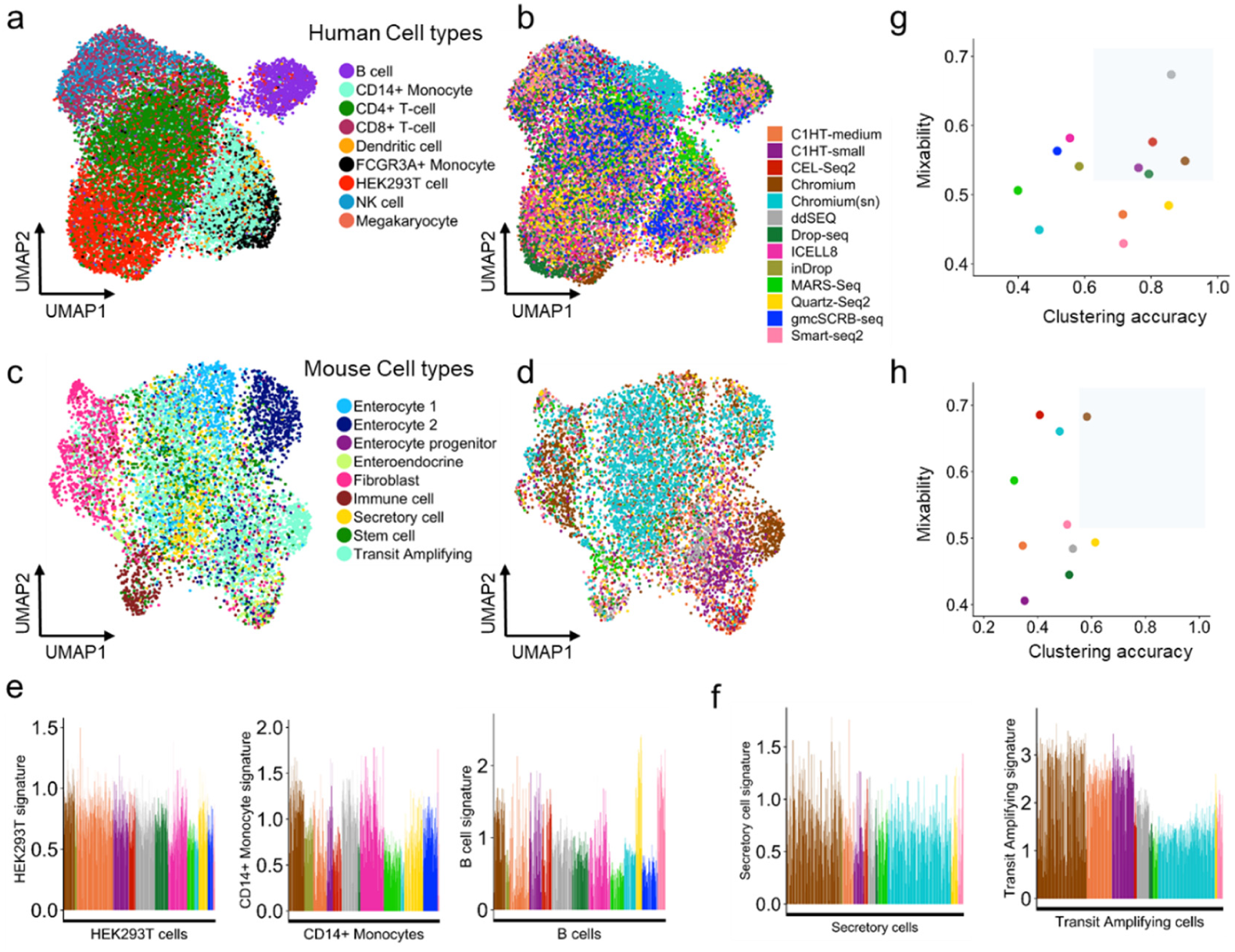
Integration of sc/snRNA-seq methods. UMAP visualization of cells after integrating technologies for human (**a,b**) and mouse (**c,d**) datasets. Cells are colored by cell type (**a,c**) and sc/snRNA-seq protocol (**b,d**). **e,f.** Barplots showing normalized and method-corrected (integrated) expression scores of cell-type-specific signatures for human HEK293T cells, monocytes, B-cells (**e**) and mouse secretory and transit-amplifying cells (**f**). Bars represent cells and colors methods. **g,h.** Evaluation of method integratability in human (**g**) and mouse (**h**). Protocols are compared according to their ability to group cell types into clusters (after integration) and mix with other technologies within the same clusters. Points are colored by sequencing method. The gray area shows the optimal trade-off between the two properties.

**Figure 6.**
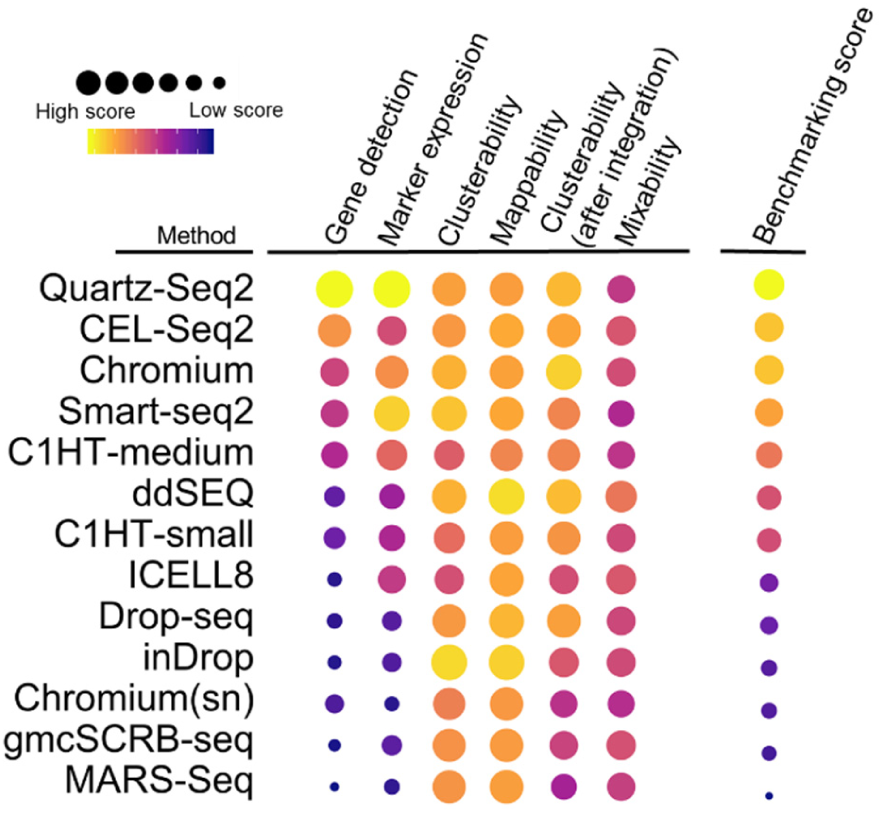
Benchmarking summary of 13 sc/snRNA-seq methods. Methods are scored by key analytical metrics, characterizing protocols according to their ability to recapitulate the original structure of complex tissues, and their suitability for cell atlas projects. The methods are ordered by their overall benchmarking score, which is computed by averaging the scores across metrics.

### Technical effects and information content

We further assessed the magnitude of technical biases, and the methods’ ability to describe cell populations. To quantify the technical variation within and across protocols, we selected highly variable genes (HVG) across all datasets, and plotted the variation in the main principle components (PC; **Figure 3a**). Using the downsampled data for HEK293T cells, monocytes and B-cells, we observed a strong protocol-specific signature, with the main source of variability being the number of genes detected per cell (**Figure 3b**). Nevertheless, PC analysis also showed a mixing of the data points for cells from different methods, suggesting generally conserved information content across the methods. Data from snRNA-seq did not show notable outliers, indicating conserved representation of the transcriptome between the cytoplasm and nucleus. The technical effects were also visible when using t-distributed stochastic neighbor embedding (tSNE) as non-linear dimensionality reduction method (**Supplementary Fig. 7**). By contrast, the methods largely mixed when the analysis was restricted to cell type-specific marker genes, suggesting a conserved cell identity profile across techniques (**Supplementary Fig. 8**).

Next, we quantified the similarities in information content of the protocols. Again, we used the downsampled datasets and calculated the correlation between methods in average transcript counts across multiple cells, thus compensating for the sparsity of single-cell transcriptome data. For the three human cell types, we observed a broad spectrum of correlation between technologies, with generally lower correlation for smaller cell types (**Figure 3c**). Here, the Chromium snRNA-seq protocol displayed a notable outlier, possibly driven by a decreased correlation of immature transcripts (intronic counts; **Figure 1**). Restricting the correlation analysis to population-specific marker genes, we observed less variation between techniques (Pearson’s r, 0.5-0.7), which underlines the fact that the expression of these markers is largely conserved between the methods (**Supplementary Fig. 9**).

To further test the suitability of protocols to describe cell types, we determined their sensitivity to detect population specific expression signatures, and found that they had remarkably variable power to detect marker genes (**Figure 3d,e**). Although most of the marker genes were detected by all technologies (>83% of genes), the magnitude of detection varied substantially. Quartz-seq2 and Smart-seq2 showed high expression levels for all cell type signatures, indicating that they have higher power for cell type identification. Since marker genes are particularly important for data interpretation (e.g. annotation), low marker detection levels could severely limit the interpretation of poorly explored tissues, or when trying to identify subtle differences between subpopulations.

The methods also detected vastly different total numbers of genes when accumulating transcript information over multiple cells, with strong positive outliers observed for the smaller cell types (**Figure 3f**). In particular, CEL-seq2 and Quartz-seq2 identified many more genes than other methods. Intriguingly, CEL-seq2 outperformed all other methods by detecting many weakly expressed genes; genes detected specifically by CEL-seq2 had significantly lower expression than the common genes detected by Quartz-seq2 (p<2.2e-16). The greater sensitivity to weakly expressed genes makes this protocol particularly suitable for describing cell populations in detail, an important prerequisite for creating a comprehensive cell atlas and functional interpretation. To further illustrate the power of the different protocols to chart the heterogeneity of complex samples, we clustered and plotted downsampled datasets in two-dimensional space (**Figure 4a**) and then calculated the cluster accuracy and Average Silhouette Width (ASW^18^, **Figure 4b**), a commonly used measure for assessing the quality of data partitioning into communities. Consistent with the assumption that library complexity and sensitive marker detection provides greater power to describe complexity, methods that performed well for these two attributes showed better separation of subpopulations, greater ASW and cluster accuracy. This is illustrated in the monocytes, for which accurate clustering protocols separated the major subpopulations (CD14+ and FCGR3A+), while methods with low ASW did not distinguish between them. Similarly, several methods were able to distinguish between CD8+ and NK cells, while others were not.

### Joint analysis across datasets

A common scenario for cell atlas projects is that data are produced at different sites using different scRNA-seq protocols. However, the final atlas is created from a combination of datasets, which requires that the technologies used are compatible. To assess how suitable it is to combine the results from our protocols into a joint analysis, we used downsampled human and mouse datasets to produce a joint quantification matrix for all techniques^19^. Importantly, single cells grouped themselves by cell type, suggesting that cell phenotypes are the main driver of heterogeneity in the joint datasets (**Figure 5a-d** and **Supplementary Fig. 10a,b**). Indeed, the combined data showed a clear separation of cell states (e.g. T-cell and enterocyte subpopulations) and rarer cell types, such as dendritic cells. However, within these populations there were some differences between the methods, indicating the presence of technical effects that could not be entirely removed during the merging step (**Figure 5e-f** and **Supplementary Fig.10c,d**). To formally assess the capacity of the methods to be joined, we calculated the degree to which technologies mix in the merged datasets (**Figure 5g,h**). Intriguingly, the methods’ suitability to be combined was not directly correlated with their power to discriminate between cell types. Thus, while a well-performing method might result in a high-resolution cell atlas maps, it could perform poorly in a consortium-driven project that includes different data sources. Moreover, when integrating further downsampled datasets, we observed a drop in mixing ability, although the cell type separation was largely conserved (**Supplementary Fig. 10e**). Consequently, quality standard guidelines for consortia might define minimum coverage thresholds to ensure the subsequent option of data merging.

Cell atlas datasets will serve as a reference for annotating cell types and states in future experiments. Therefore, we assessed cells’ ability to be projected onto our reference sample (**Figure 2b,c**). We used the population signature model defined by *matchSCore2* and evaluated the protocols based on their cell-by-cell mapping probability, which reflects the confidence of cell annotation (**Supplementary Fig. 11a-c**). Although there were some differences in the protocols’ projection probabilities and a potential bias due to the selection of the reference protocol, a confident annotation was observed for most cells with inDrop and ddSEQ reporting the highest probabilities. Notably, high probability scores were also observed in further downsampled datasets (**Supplementary Fig. 11b**). This has practical consequences, as data derived from less well performing methods (from a cell atlas perspective) or from poorly sequenced experiments could be identifiable and thus suitable for specific analysis types, such as tissue composition profiling.

## Conclusion

Systematic benchmarking of available technologies is a crucial prerequisite for large-scale projects. Here, we evaluated scRNA-seq protocols for their power to produce a cellular map of complex tissues. Our reference sample simulated common scenarios in cell atlas projects, including differentiated cell types and dynamic cell states. We defined the strengths and weaknesses of key features that are relevant for cell atlas studies, such as comprehensiveness, integratability, and predictive value. The methods revealed a broad spectrum of performance, which should be considered when defining guidelines and standards for international consortia (**Figure 6**). In addition, cell atlas projects need to consider other protocol-specific features, such as cost-effectiveness and scalability, in their decision making process towards large-scale datasets. It is equally important to benchmark computational pipelines for data analysis and interpretation^20–22^. We envision that the datasets provided by our study will serve as a valuable resource for the single-cell community to develop and evaluate novel strategies towards an informative and interpretable cell atlas.

## Addendum

This study complements a study entitled “Systematic comparative analysis of single cell RNA-sequencing method” by Ding et al., which applied a complementary design (BIORXIV/2019/ 632216).

## Acknowledgement

This project has been made possible in part by grant number 2018-182827 from the Chan Zuckerberg Initiative DAF, an advised fund of Silicon Valley Community Foundation. HH is a Miguel Servet (CP14/00229) researcher funded by the Spanish Institute of Health Carlos III (ISCIII). CM is supported by an AECC postdoctoral fellowship. This work has received funding from the European Union’s Horizon 2020 research and innovation programme under Marie Skłodowska-Curie grant agreement No. H2020-MSCA-ITN-2015-675752 (Singek), and the Ministerio de Ciencia, Innovación y Universidades (SAF2017-89109-P; AEI/FEDER, UE). S. is supported by the DFG (GR4980) the Behrens-Weise-Foundation. D.G. and S. are supported by the Max Planck Society. CZ is supported by EMBO through long-term fellowship ALTF 673-2017. The single-nucleus RNA-Seq data were generated with support from the National Institute of Allergy and Infectious Diseases grant U24AI118672, the Manton Foundation, and the Klarman Cell Observatory (AR). We thank ThePaperMill for critical reading and scientific editing services and the Eukaryotic Single Cell Genomics Facility at Scilifelab (Stockholm, Sweden) for support. Core funding is from the ISCIII and the Generalitat de Catalunya.

## Ethical Statement

This study was approved by the Parc de Salut MAR Research Ethics Committee (reference number: 2017/7585/I) to Dr. Holger Heyn. We adhered to ethical and legal protection guidelines for human subjects, including informed consent.

## Author’s Contributions

HH designed the study. EM and AL performed all data analyses. CM, AA and EB prepared the reference sample. CZ, DJM, SP and OS supported the data analysis. MG and IG provided technical and sequencing support. All other authors provided sequencing-ready single-cell libraries or sequencing raw data. HH, EM and AL wrote the manuscript with contributions from the co-authors. All authors read and approved the final manuscript.

## Data Availability

All raw sequencing data will be freely available through the Human Cell Atlas Data Coordination Portal (DCP). The code for *matchSCore2* is available under https://github.com/elimereu/matchSCore2.

## Online Methods

### Reference sample

#### Cell Lines

NIH3T3-GFP, MDCK-TurboFP650 and HEK293-RFP were cultured at 37°C in an atmosphere of 5% (v/v) carbon dioxide in Dulbecco’s Modified Eagle’s Medium, supplemented with 10% (w/v) fetal bovine serum, 100 U penicillin, and 100 µg/L streptomycin (Invitrogen). On the reference sample preparation day, the culture medium was removed and the cells washed with 1X PBS. Afterwards, cells were trypsinized (trypsin 100X), pelleted at 800 × g for 5min, washed in 1X PBS, re-suspended in PBS-EDTA (2mM) and stored on ice.

#### Mouse Colon Tissue

The colon from eleven mice (7x*LGR5/GFP* and 4WT) was dissected and removed. For single-cell separation the colons were treated separately. The colon was sliced, opened and washed twice in cold 1X HBSS. It was then placed on a petri plate on ice and minced with razor blades until disintegration. The minced tissue was transferred to a 15 ml tube containing 5 ml of 1X HBSS and 83 µl of collagenase IV (final concentration 166 U/ml). The solution was incubated for 15 min at 37°C (vortexed for 10 sec every 5 min). To inactivate the collagenase IV, 1 ml of FBS was added and vortexed for 10 seconds. The solution was filtered through a 70 µm nylon mesh (changed when clogged). Finally, all samples were combined, cells pelleted for 5 min at 400 g at 4°C. The supernatant was removed, and the cells resuspended in 20 ml of 1X HBSS and stored on ice.

#### Isolation of peripheral blood mononuclear cells (PBMC)

Whole blood was obtained from four donors (two female, two male). The extracted blood was collected in Heparin tubes (GP supplies) and processed immediately. For each donor, PBMCs were isolated according to the manufacturer’s instructions for FICOLL extraction (pluriSelect). Briefly, blood from two Heparin tubes (approximately 8 ml) was combined, diluted in 1X PBS and carefully added to a 50 ml tube containing 15 ml FICOLL. The tubes were centrifuged for 30 min at 500 g (minimum acceleration and deceleration). The interphase was carefully collected and diluted with 1X PBS + EDTA (2mM). Following a second centrifugation, the supernatant was discarded and the pellet resuspended in 2 ml of 1X PBS + EDTA (2mM) and stored on ice.

#### Preparation of the reference sample

Cell counting was performed using an automated cell counter (TC20™ Automated Cell Counter, Bio-Rad Laboratories). The reference sample was calculated to include human PBMC (60%), mouse colon (30%), HEK293T (6%, RFP labelled human cell line), NIH3T3 (3%, GFP labelled mouse cells) and MDCK (1%, TurboFP650 labelled dog cells). To adjust for cell integrity loss during sample processing, we measured the viability during cell counting, and accounted for an expected viability loss after cryopreservation (10% for cell lines and PBMC; 50% for colon^17^). All single cell solutions were combined in the proportions mentioned above and diluted to 250,000 viable cells per 0.5 ml. For cryopreservation, 0.5 ml of cell suspension was aliquoted into cryotubes and gently mixed with a freezing solution (final concentration 10% DMSO; 10% heat-inactivated FBS). Cells were then frozen by gradually decreasing the temperature (1°C/min) to – 80°C (cryopreserved), and stored in liquid nitrogen. MARS-Seq and Smart-Seq2 experiments were performed to validate sample quality and composition before distributing aliquots to the partners.

#### Sample processing instructions

This cryopreserved reference sample forms the basis for systematic comparison of scRNA-seq techniques. The sample consists of two complex tissues (human PBMC and mouse colon) and three cell lines (HEK293-RFP, NIH3T3-GFP, MDCK-Turbo650). The primary PBMC and the colon cells account for around 90% of the living (DAPI negative) sample content, and the cell lines the remaining 10% (6% HEK293-RFP, 3% NIH3T3-GFP, <1% MDCK-Turbo650). Each cryo-vial contains ∼250,000 living cells, sufficient to sort a minimum of 4 × 384-well plates or to isolate >3000 cells (microfluidic systems), and should be stored at −80°C upon arrival. The sample preparation aims to be standardized for all methods to allow comparison of the performance of library preparation. FACS isolation should be performed before sample processing to exclude damaged/dead cells (DAPI positive). Moreover, we aim to simulate the exclusion of unwanted cell types by excluding NIH3T3 (GFP positive) cells during FACS isolation. For the remaining sample, FACS gates should be set to exclude debris, cell fragments and doublets (Appendix: screen shots provided). Proportions of intact (DAPI negative) and fluorescence labeled cells (RFP, GFP and TurboFP650) should be recorded, and, if possible, cells should be index-sorted (for microtiter plates).

**NOTE:** The cryopreserved samples consists of approximately 30-40% intact (DAPI negative) cells. We recommend FACS isolation of DAPI negative cells before single-cell capture.

**NOTE:** We provide cryo-vials of PBMC and fluorescence-labeled cell lines to facilitate gate-setting for debris exclusion, and to define the degree of compensation. Please set gates to include blood and larger cells as indicated in the Appendixes.

**NOTE:** One HCA reference vial is sufficient to fill 4x 384-well plates.

**NOTE:** FACS isolation into plates should be performed at low speed (below 100 cells/sec) to avoid loss of the sample.

**NOTE:** To simulate the exclusion of cell types, GFP-labeled NIH3T3 cells should be excluded from the final single-cell selection.

#### Sample thawing instructions

- Remove sample from −80°C and process immediately
- De-freeze in water bath (37°C) with continuous agitation until material is almost thawed
- Transfer to 15 ml Falcon using a 1000 ul tip (wide-bored or cut tip) without mixing by pipetting
- Add drop-wise 1000 ul of pre-warmed (37°C) Hibernate-A while gently swirling the sample
- Let sample rest for 1 min
- Add drop-wise 2000 ul of pre-warmed (37°C) Hibernate-A while gently swirling the sample
- Let sample rest for 1 min
- Add drop-wise 2000 ul of pre-warmed (37°C) Hibernate-A while gently swirling the sample
- Let sample rest for 1 min
- Add 3000 ul pre-warmed (37°C) Hibernate-A
- Invert Falcon 3 times
- Let sample rest for 1 min
- Add 5000 ul pre-warmed (37°C) Hibernate-A
- Invert Falcon 6 times
- Let sample rest for 1 min
- Centrifuge sample at 400 g for 5 min at 4°C (pellet clearly visible)
- Remove supernatant until 500 ul supernatant remains in tube
- Resuspend the pellet by gentle pipetting
- Add 3500 ul of 1X PBS + 2mM EDTA
- Store sample on ice until processing
- Filter cells through a nylon mesh into FACS tubes (2 tubes with 2 ml sample)
- Add 3 ul DAPI
- Mix gently
- Store on ice
- **Exclude DAPI and GFP positive cells during sorting**
- Use index sorting for RFP and TurboFP650 (optional)

### Single-cell RNA sequencing library preparation

#### Quartz-Seq2^23^

We isolated single-cells into 384-well PCR plates from cell suspension using a MoFlo Astrios EQ (Beckman Coulter) cell sorter. The cell sorter was equipped with a 100-μm nozzle and a custom-made splash-guard. In total, we analyzed 3,072 wells corresponding to eight 384-well PCR plates. Sequence library preparation of Quartz-Seq2 was performed as described previously^23^ with the following modifications. For lysis buffer, we used 768 kinds of RT primers corresponding to v3.2A and v3.2B. We prepared two sets of the 384-well PCR plate with lysis buffer containing no ERCC spike-in RNA. We added 1 μl of RT premix (2X Thermopol buffer, 1.25 units/μL SuperScript III, 0.1375 units/μL RNasin plus) to 1 μl of lysis buffer for each well. After cell barcoding, we collected cDNA solution into one well reservoir from two sets of 384-well plates, which corresponded to 768 wells. For cDNA purification and concentration, we used four purification columns for cDNA solution from two 384-well PCR plates. In the PCR step, we amplified cDNA for ten cycles. In an additional purification step for amplified cDNA, we added 26 μl (0.65X) of resuspended AMPure XP Beads to the cDNA solution. We obtained amplified cDNA of 32.6 ± 6.8 ng (n = 4) from the 768 wells. We sequenced the Quartz-Seq2 sequence library with a NextSeq 500/550 High Output v2 Kit. The BCL files obtained were converted to FASTQ files using bcl2fastq2 (v2.17.1.14) with demultiplexing pool barcodes. Each FASTQ file was split into single FASTQ files for each cell barcode using a custom script (https://github.com/rikenbit/demultiplexer_quartz-seq2, DOI: 10.5281/zenodo.2585429).

#### inDrop System (1CellBio)^24^

Cells were isolated using an Aria3Fusion (BD Bioscience) cell sorter with a 100µm nozzle and a flow rate of 6-7. The workflow was carried out using the inDrop instrument and the inDrop single cell RNA-seq kit (Cat. No. 20196, 1CellBio) according to the manufacturer’s protocols. Microfluidic chips were prepared by silanization, and barcode labeled hydrogel microspheres (BHMs) were prepared shortly before cell capture, according to protocol (version v2.0., 1CellBio website). Droplet-making oil, single-cell suspension (200 cells/µL), and freshly prepared RT/lysis buffer were loaded onto the chip for droplet generation, according to the inDrop protocol for single-cell encapsulation and reverse transcription (version 2.1., 1CellBio website). An emulsion corresponding to ∼4000 droplets was collected in a cooled tube and irradiated with UV light to release the photo-cleavable barcoding oligos from the BHMs. cDNA synthesis proceeded within the droplets, and the emulsion was subsequently split into equal volumes in such a way as to not exceed ∼2000 droplets per reaction tube. After de-emulsification, cDNA contained in the aqueous phase was stored at −80°C. The RT product was further processed according to the InDrop library preparation protocol (version 1.2. 1CellBio website). The cDNA was fragmented by ExoI/HinfI and purified by AMPure XP beads. Second strand synthesis was conducted using NEB second-strand synthesis module (Cat. no. E6111S, NEB). In vitro-transcription was conducted using HiScribe T7 High Yield RNA Synthesis kit (cat. no. E2040S, NEB). Amplified RNA was then fragmented, and the fragments used in a second reverse transcription reaction with random hexamers to convert the sample back into DNA and to add a read primer-binding site to each molecule. Hybrid molecules of RNA and DNA were cleaned up using AMPure beads and amplified by PCR. Final libraries were sequenced using HiSeq4000 and NextSeq (Illumina).

#### ICELL8 SMARTer Single-Cell System (Takara Bio)^25^

Hoechst 33342 and propidium iodide co-stained single-cell suspension (20 cells/µL) was distributed in eight wells of a 384-well source plate (Cat. No. 640018, Takara) and dispensed into a barcoded SMARTer ICELL8 3’ DE Chip (Cat. No. 640143, Takara) using an ICELL8 MultiSample NanoDispenser (MSND, Takara). 4 chips were used to target ∼3000 single cells. Nanowells were imaged using the ICELL8 Imaging Station (Takara). After imaging, the chip was sealed, placed in a pre-cooled freezing chamber, and stored at −80 °C. CellSelect software was used to identify each nanowell that contained a single cell. These nanowells were then selected for subsequent targeted deposition of a 50 nL/nanowell RT-PCR reaction solution from the SMARTer ICELL8 3’ DE Reagent Kit (Cat. No. 640167, Takara) using the MSND. After RT and amplification in a Chip Cycler, barcoded cDNA products from nanowells were pooled together using the SMARTer ICELL8 Collection Kit (Cat. No. 640048, Takara). cDNA was concentrated using the Zymo DNA Clean & Concentrator kit (Cat. No. D4013, Zymo Research), and purified using AMPure XP beads. cDNA was then used to construct Nextera XT (Illumina) DNA libraries, followed by 0.6X AMPure XP bead purification. Library quantification and size distribution was done using Qubit, KAPA Library Quantification and Agilent TapeStation. Final libraries were sequenced using HiSeq4000 and NextSeq500 (Illumina).

#### Drop-Seq (Dolomite)^26^

Single-cell RNA experiments were performed using the scRNA system with P-Pumps and a scRNA-chip (100µm channel width) from Dolomite Bio (Royston, UK). Encapsulation was conducted according to the manufacturer’s instructions, and library construction was completed according to the published DropSeq protocol^26^. Briefly, polyT-barcoded beads (MACOSKO-2011-10; ChemGenes) were loaded at a concentration of 600/µl, and cells at a concentration of 450/µl. The pumps were operated at a flowrate of 30 µl/min for beads and cell suspension (PBS+2mM EDTA), and at 200 µl/min for oil (QX200™ Droplet Generation Oil for EvaGreen; BioRad). After encapsulation, cell lysis, and hybridization of RNA to the beads, droplets were broken using PFO (Sigma-Aldrich) and aliquots of a maximum of 90000 beads were collected. Reverse transcription was performed in a 200µl volume with Maxima H Minus Reverse Transcriptase (Thermo Fisher Scientific) and 2.5 µM TSO-primer (AAGCAGTGGTATCAACGCAGAGTGAATrGrGrG; Qiagen) at room temperature for 30 min, followed by 90 min at 42°C. After exonuclease treatment (ExoI; New England Biolabs) at 47°C in 200 µl, to digest the unbound primer, cDNA was amplified by PCR using HiFi HotStart mix (Kapa Biosystems) and amplification primer (AAGCAGTGGTATCAACGCAGAGT; Qiagen) in batches of 4000 beads in a volume of 50 µl (95°C – 3min; 4 cycles: 98°C – 20s, 65°C – 45s, 72°C – 3min; 9 cycles: 98°C – 20s, 67°C – 20s, 72°C – 3min; 72°C – 5min). Libraries were generated using the Nextera XT library Kit (Illumina) in five pooled PCR samples with 600 pg of cDNA and a custom P5-primer (AATGATACGGCGACCACCGAGATCTACACGCCTGTCCGCGGAAGCAGTTGGTATC AACGCAGAGT*A*C; Qiagen). Final library QC was conducted using the BioAnalyzer High Sensitivity DNA Chip (Agilent Technologies). For sequencing on an Illumina HiSeq2500 V4, we used a custom read 1 primer (GCCTGTCCGCGGAAGCAGTGGTATCAACGCAGAGTAC; Qiagen).

#### Chromium V2 (10X Genomics): Single-cell RNA sequencing^15^

Two cell preparations were conducted on two different days: one to prepare 2 libraries for sequencing at high read depth, and one to prepare 8 libraries at low read depth. To prepare the libraries for high read depth, one frozen vial of a Human Cell Atlas reference sample was thawed and prepared as described. At the end of this protocol, the cells were resuspended in PBS with 2 mM EDTA. Since cells showed clumping and low viability, they were centrifuged 3 times at 150 g for 10 min at room temperature, and resuspended in 50% PBS, 2mM EDTA and 50% Iscove’s Modified Dulbecco Medium (IMDM, ATCC) supplemented with 10% FBS and filtered through a 40µm FlowMi cell strainer (Sigma-Aldrich) to remove cell aggregates and large cell debris. At the final count before loading, the cell suspension demonstrated a viability of 60%. To prepare the libraries for low read depth, two frozen vials of a the reference sample were thawed and prepared as described in an updated version of the HCA Benchmark protocol. At the end of this protocol, the cells were resuspended in IMDM, 10% FBS and 1mM EDTA, and filtered through a 40-µm FlowMi cell strainer to remove cell aggregates and large cell debris. At the final count before loading, the cell suspension demonstrated a viability of 65%. The cells were not processed using FACS isolation, but run directly on the 10x Chromium system (10x Genomics, Pleasanton, CA, USA). Cells were mixed with single-cell master mix, and the resulting cell suspensions were loaded on a 10x Chromium system to generate 2 libraries at 5,000 cells each and 5 libraries at 10,000 cells each. The single-cell libraries were generated using 10x Chromium Single Cell gene expression V2 reagent kits according to the manufacturer’s instructions (Chromium single cell 3’ reagents kits v2 user guide). Single cell 3’ RNA-seq libraries were quantified using an Agilent Bioanalyzer with a high sensitivity chip (Agilent), and a Kapa DNA quantification kit for Illumina platforms (Kapa Biosystems). The libraries were pooled according to the target cell number loaded. Sequencing libraries were loaded at 200 pM on an Illumina NovaSeq6000 with Novaseq S2 Reagent Kit (100 cycles) using the following read lengths: 26 bp Read1, 8 bp I7 Index and 91 bp Read2. The 2 libraries of 5,000 cells and the 8 libraries of 10,000 cells were sequenced at 250,000 and 25,000 reads per cell, respectively.

#### Chromium V2 (10X Genomics): Single-nucleus RNA sequencing

We isolated nuclei from the cell suspension using a protocol provided by 10x Genomics^27^. We counted the nuclei using a Countess II (Thermo Fisher Scientific). We made an aliquot containing ∼11,000 nuclei in a volume of 33.8 µL in RB buffer (1x PBS, 1% BSA, and 0.2U/µl RNaseIn (TaKaRa)) as sample A, and stained the rest of the nuclei suspension with Vybrant DyeCycle Violet Stain (Thermo Fisher Scientific) at a concentration of 10 µM. We used a MoFlo Astrios EQ cell sorter (Beckman Coulter) and set fluorescence activated cell sorting (FACS) gating on forward scatter plot, side scatter plot and on fluorescent channels to pick Violet-positive (for nuclei), while excluding debris and doublets. We used a 100 µm nozzle to sort 20,000 nuclei into 20 μl RB buffer as sample B. After sorting, we measured the volume of B with a pipette, spun it at 500 g for 5 min at 4°C, and then carefully removed part of the supernatant to leave ∼40µl. We resuspended B by gentle pipetting 40 times.

Immediately after nuclei isolation, we loaded sample A into one channel of a Chromium Single Cell 3’ Chip (10x Genomics, PN-120236), and then processed it through the Chromium Controller to generate GEMs (Gel Beads in Emulsion). We then loaded 33.8 µL of B 25 minutes later after sorting and centrifugation, as described above, into one channel of a second chip, and processed it in the same way as the first chip. We prepared RNA-Seq libraries for both samples in parallel with the Chromium Single Cell 3’ Library & Gel Bead Kit V2 (10x Genomics, PN-120237), according to the manufacturer’s protocol. We pooled the 2 samples based on molar concentrations and sequenced them on a NextSeq500 instrument (Illumina).

#### Smart-seq2^28^

Smart-seq2 libraries were prepared at half the volume, as described previously^28^, with minor modifications. In brief, 2 µl of lysis buffer containing 0.1 % Triton X-100 (Sigma-Aldrich), 1 U/µl RNase inhibitor (Takara), 2.5 mM dNTPs (Thermo Fisher) and 2 µM oligo-dT primer (5′– AAGCAGTGGTATCAACGCAGAGTACT30VN-3′; IDT) were dispensed into each well of a 384-well plate (4titude). Lysis plates were stored at −20°C until cell sorting, after which single-cell lysates were kept at −80 °C. Before reverse transcription, cell lysates were denatured at 72 °C for 3 min and immediately placed on ice. The RT reaction was performed in a 5 µl total volume, with final reagent concentrations of 1x Superscript first-strand buffer (Thermo Fisher), 5 mM DTT (Thermo Fisher), 1 M Betaine (Sigma-Aldrich), 9 mM MgCl2 (Sigma-Aldrich), 1 U/µl RNase inhibitor (Takara), 1 µM LNA template-switching oligo (5′-AAGCAGTGGTATCAACGCAGAGTACATrGrG+G-3′; Exiqon), and 10 U/µl Superscript II RT enzyme. Next, pre-amplification PCR was performed for 22 cycles at final concentrations of 1x KAPA HiFi HotStart ReadyMix (Roche) and 0.08 µM ISPCR primer (5′-AAGCAGTGGTATCAACGCAGAGT-3′; IDT) in a total reaction volume of 11 µl. The cDNA was cleaned up by adding 10 µl of SPRI beads (19.5 % PEG, 1 M NaCl, 1 mM EDTA, 0.01 % IGEPAL CA-630), washing twice with 20 µl 80 % ethanol, and eluting in 10 µl H2O. The cDNA concentration was measured for all wells using Picogreen dsDNA assay (Thermo Fisher), and diluted to 200 pg/µl using a Mantis liquid handler (Formulatrix). Next, 1 µl of cDNA was used as input for the Nextera XT library preparation kit (Illumina) at 1/5 volume, according to the manufacturer’s instructions. During the 12 cycles library PCR, custom i7 and i5 indexing primers (IDT) were added at 0.5 µM each. Finally, 5 µl of library per well were pooled, cleaned and concentrated using SPRI beads (19.5 % PEG; see above). Final libraries were sequenced using HiSeq2500 V4 (Illumina).

#### CEL-Seq2^29,30^

Single-cell RNA sequencing was performed using a modified version of the mCEL-Seq2 protocol, an automated and miniaturized version of CEL-Seq2, on a Mosquito nanoliter-scale liquid-handling robot (TTP LabTech)^29,31^. Briefly, cells were sorted into 384-well plates (Bio-Rad) containing 240 nl of lysis buffer containing polyT primers and 1.2 μl of mineral oil (Sigma-Aldrich). Sorted plates were centrifuged at 2200 g for several minutes at 4°C, snap-frozen in liquid nitrogen and stored at −80°C until processing. 160nL of reverse transcription reaction mix and 2.2 μl of second strand reaction mix were used to convert RNA into cDNA. cDNA from 96-cells were pooled together before clean up and *in vitro* transcription, generating 4 libraries from one 384-well plate. During all purification steps, including the library cleanup, we used 0.8 μl of AMPure/RNAClean XP beads (Beckman Coulter) per 1 μl of sample. Sixteen libraries with 96 cells each (one of the libraries contained 30,000 RNA molecules from ERCC spike-in mix per cell) were sequenced on an Illumina HiSeq3000 sequencing system (pair-end multiplexing run).

#### MARS-Seq^32^

To construct single-cell libraries from poly(A)-tailed RNA, we used massively parallel single-cell RNA sequencing (MARS-Seq)^32^. Briefly, single cells were FACS-isolated into 384-well plates containing lysis buffer (0.2% Triton X-100 (Sigma-Aldrich); RNase inhibitor (Invitrogen)) and reverse-transcription (RT) primers. Single-cell lysates were denatured and immediately placed on ice. The RT reaction mix, containing SuperScript III reverse transcriptase (Invitrogen), was added to each sample. After RT, the cDNA was pooled using an automated pipeline (epMotion, Eppendorf). Unbound primers were eliminated by incubating the cDNA with exonuclease I (NEB). A second stage of pooling was performed through cleanup with SPRI magnetic beads (Beckman Coulter). Subsequently, pooled cDNAs were converted into double-stranded DNA using the Second Strand Synthesis enzyme (NEB), followed by clean-up and linear amplification by T7 *in vitro* transcription overnight. The DNA template was then removed by Turbo DNase I (Ambion), and the RNA purified using SPRI beads. Amplified RNA was chemically fragmented using Zn2+ (Ambion), and then purified using SPRI beads. The fragmented RNA was ligated with ligation primers containing a pool barcode and partial Illumina Read1 sequencing adapter using T4 RNA ligase I (NEB). The ligated products were reverse-transcribed using the Affinity Script RT enzyme (Agilent Technologies) and a primer complementary to the ligated adapter, partial Read1. The cDNA was purified using SPRI beads. Libraries were completed by a PCR step using the KAPA Hifi Hotstart ReadyMix (Kapa Biosystems) and a forward primer containing the Illumina P5-Read1 sequence, and a reverse primer containing the P7-Read2 sequence. The final library was purified using SPRI beads to remove excess primers. Library concentration and molecular size were determined with a High Sensitivity DNA Chip (Agilent Technologies). Multiplexed pools were run on Illumina HiSeq2500 Rapid flow cells (Illumina).

#### C1 High-Throughput (HT-IFC)^33^

Cells were sorted into 15-ml tubes containing 7 ml of PBS with 5% FBS, using a Sony SH800 Cell Sorter. Cells were concentrated by centrifugation at 350 g for 5 minutes at 4°C. The supernatant was removed, and cells were counted and diluted to 900 cells/ul for the Fluidigm C1 HT Small-Cell Integrated Fluidic Circuits (IFCs), and 450 cells/ul for the Fluidigm C1 HT Medium-Cell IFCs. A total of eight small-cell and seven medium-cell IFCs were used to generate cDNA on the Fluidigm C1 System. cDNA generation and the subsequent preparation of sequencing libraries were performed according to the recommended Fluidigm C1 HT protocols. Enrichment Primers from the Fluidigm reagent kit were replaced with NEBNext i5xx primers from NEBNext Multiplex Oligos for Illumina (Dual Index Primers Set 1 & 2) (New England BioLabs), to enable library pooling. Libraries from fifteen IFCs were pooled and sequenced on the NovaSeq6000 system (Illumina) in two runs on the S2 flow cell.

#### ddSEQ (Bio-Rad)

Flow cytometry analysis and cell sorting were performed on the S3e Cell Sorter using ProSort Software (Bio-Rad Laboratories, #12007058) for acquisition and sorting. Viable cells were sorted into 1x PBS with + 0.1% BSA and kept at 4°C until scRNA-Seq. Cell concentration of sorted cells was determined using the TC20 Automated Cell Counter (Bio-Rad Laboratories, #1450102) and adjusted to a final concentration of 2,500 cells/ul. Cells were then prepared for single-cell sequencing using the Illumina Bio-Rad SureCell WTA 3’ Library Prep Kit for the ddSEQ (Illumuina, #20014280). Cells were loaded onto ddSEQ cartridges and processed in the ddSEQ Single-Cell Isolator (Bio-Rad Laboratories, #12004336) to isolate and barcode single cells in droplets. First-strand cDNA synthesis occurred in droplets, which were then disrupted for second strand cDNA synthesis in bulk. Libraries were prepared according to manufacturer’s instructions and then sequenced on the NextSeq500 system (Illumina).

#### gmcSCRB-seq^34^

Cells were sorted and processed using the alternative lysis (Guanidin) condition (gmcSCRB-seq) as described suitable for PBMCs in Bagnoli et al (2018). Briefly, single cells (“3 drops” purity mode) were sorted into 96-well DNA LoBind plates (Eppendorf) containing 5 µl lysis buffer using a Sony SH800 sorter (Sony Biotechnology; 100 µm chip). Lysis buffer consisted of 5 M guanidine hydrochloride (Sigma-Aldrich), 1% 2-mercaptoethanol (Sigma-Aldrich) and a 1:500 dilution of Phusion HF buffer (New England Biolabs). Samples were processed in six batches, with one batch of two plates and five batches of six plates. Each well was cleaned up using SPRI beads and resuspended in 4 µl H2O (Invitrogen) and a mix of 5 µl reverse transcription master mix, consisting of 20 units Maxima H- enzyme (Thermo Fisher), 2 × Maxima H- Buffer (Thermo Fisher), 2 mM each dNTPs (Thermo Fisher), 4 µM template-switching oligo (IDT), and 15% PEG 8000 (Sigma-Aldrich). For libraries containing ERCCs, 30,000 molecules of ERCC spike-in Mix 1 (Ambion) was used and the H2O (Invitrogen) was adjusted accordingly. After the addition of 1 µl 2 µM barcoded oligo-dT primer (E3V6NEXT, IDT), cDNA synthesis and template switching was performed for 90 min at 42 °C. Barcoded cDNA was then pooled in 2 ml DNA LoBind tubes (Eppendorf) and cleaned up using SPRI beads. Purified cDNA was eluted in 17 µl and residual primers digested with Exonuclease I (Thermo Fisher) for 20 min at 37 °C. After heat inactivation for 10 min at 80 °C, 30 µl PCR master mix consisting of 1.25 U Terra direct polymerase (Clontech) 1.66 × Terra direct buffer and 0.33 µM SINGV6 primer (IDT) was added. PCR was cycled as given: 3 min at 98 °C for initial denaturation followed by 19 cycles of 15 s at 98 °C, 30 s at 65 °C, 4 min at 68 °C. Final elongation was performed for 10 min at 72 °C. Batch 4 was erroneously denatured for 10 min due to a cycler error, but left in as we consider such errors as possible batch variation errors.

Following pre-amplification, all samples were purified using SPRI beads at a ratio of 1:0.8 with a final elution in 10 µl of H2O (Invitrogen). The cDNA was then quantified using the Quant-iT PicoGreen dsDNA Assay Kit (Thermo Fisher). Size distributions were checked using high-sensitivity DNA Fragment Analyzer kits (AATI) and high-sensitivity DNA Bioanalyzer kits (Agilent). As the samples had large primer peaks, they were purified a second time using SPRI beads at a ratio of 1:0.8 and then pre-amplified for an additional 3 cycles, as above. The cDNA was then purified and reanalyzed as above. Samples passing the quantity and quality controls were used to construct Nextera XT libraries from 0.8 ng of pre-amplified cDNA. During library PCR, 3′ ends were enriched with a custom P5 primer (P5NEXTPT5, IDT). Libraries were pooled and size-selected using 2% E-Gel Agarose EX Gels (Life Technologies), cut out in the range of 300–800 bp, and extracted using the MinElute Kit (Qiagen) according to manufacturer’s recommendations. Libraries were paired-end sequenced on high output flow cells of an Illumina HiSeq 1500 instrument. Sixteen bases were sequenced with the first read to obtain cellular and molecular barcodes and 50 bases were sequenced in the second read into the cDNA fragment. An additional 8 base i7 barcode read was done to allow multiplexing.

## Data analysis

### Primary data preprocessing

FASTQ files for each technique were collected and processed in a unified manner. We developed a snakemake^35^ workflow that streamlines all steps, including read filtering and mapping, quantification, downsampling and species deconvolution, and provides a Single Cell Experiment Object^36^ output with detailed metadata. We used zUMIs^37^, a single-cell processing tool compatible with all major scRNA-Seq protocols for filtering, mapping and quantification, ensuring comparable primary data processing between all methods. First, we discarded low-quality reads (barcodes and UMI sequences with more than 1 base below the Phred quality threshold of 20) and removed barcodes with less than 100 reads.

For techniques with known barcodes, we provided zUMIs with these barcode sequences, and used the automatic barcode detection function to detect the sequenced cells for other techniques. Next, cDNA reads were mapped to the human GRCh38, mouse GRCm38, and a human-mouse-dog mixed (for species level doublet detection) reference genomes using STAR^38^. Reads were then assigned to exonic and intronic features using featureCounts^39^ and counted using the default parameters of zUMIs for human-only, mouse-only and mixed bam-files, separately. The output expression matrix of reads mapping to both exonic and intronic regions was selected for the downstream analysis. Of note, we included intronic counts in the expression quantification to improve gene detection and to enable a comparison with the snRNA-seq derived dataset. To deconvolute species, detect doublets and low quality cells, the mixed-species mapped data was used. Cells for which >70% of the reads mapped to only one species were assigned to the corresponding species. The remaining cells (those for which <70% of the reads mapped to only one species) were removed from the downstream analysis. Finally, for each technique, a *human* and *mouse* Single Cell Experiment object was created by combining the expression matrix and the metadata.

For subsequent data analysis, we discarded cells with <10,000 total number of reads as well as the cells having <65% of the reads mapped to their reference genome. Cells in the 95th percentile of the number of genes/cell and those having <25% mitochondrial gene content were included in the downstream analyses. Genes that were expressed in less than five cells were removed.

### Clustering

Filtering, normalization, selection of highly variable genes (HVG), and clustering of cells were performed using the Seurat^40^ package (version 2.3.4). The read counts were log-normalized for each cell using the natural logarithm of 1 + counts multiplied by a scale factor (10,000). To avoid spurious correlations, the library sizes were regressed out, and the genes were scaled and centered. The scaled Z-score values were then used as normalized gene measurement input for clustering and for visualizing differences in expression between cell clusters. We selected HVGs by evaluating the relationship between gene dispersion (y.cutoff = 0.5) and the log mean expression. The clustering procedure projects cells onto a reduced dimensional space, and then groups them into subpopulations by computing a shared-nearest-neighbour (SNN) based on the Euclidean distance (finding highly interconnected communities). The algorithm is a variant of the Louvain method, which uses a resolution parameter to determine the number of clusters.

In this step, the dimension of the subspace was set to the number of significant principal components (PC) based on the distribution of the PC standard deviations and by inspecting the ElbowPlot graph. The number of clusters was aligned to the expected biological variability, and cluster identities were assigned using previously described gene markers. T-SNE and UMAP were used to visualize the clustering distribution of cells. Cluster-specific markers were then identified using the Wilcoxon rank-sum test.

Trajectory analysis and pseudo-ordering of cells was performed using the Monocle^41^ package (version 2.8.0) with the previously identified HVGs. Monocle works with the raw data and allows to specify the family distribution of gene measurements, which was set to a negative binomial, as defined in the family function from the VGAM package. As for the clustering, the expression space was reduced before ordering cells using the DDRTree algorithm. To validate cell populations, and for cell type identification and annotation, we used pseudotime ordering of single cells derived from the mouse colon.

### Sample deconvolution and annotation

To identify and annotate cell types and states, we analyzed the individual single-cell experiments separately, taking advantage of the original sequencing depth. Gene expression counts were log-normalized to identify HVGs, as input to compute cell-to-cell distances and graph-based clustering (see Clustering). Cell clusters were visualized in two-dimensional space using t-SNE and UMAP, and then annotated by examining previously described cell population marker genes. All methods were able to recapitulate most cell types in both human and mouse samples, although in different proportions and resolutions.

In human samples, the T-cell marker CD3 was used to differentiate T-cells from other populations. While the CD4 T-cells cluster was clearly identifiable (with non-overlapping expression of markers), CD8 T-cells and Natural Killer (NK) were often intermixed. Monocytes were the second most abundant cell type, including subpopulations of CD14 and FCGR3A monocytes. High levels of CD79A and CD79B allowed the clear identification of B-cells. HEK293T cells generally fell into the same cluster, separate from blood subpopulations. They were clearly identifiable by the high number of detected genes (up to six-fold higher than PBMC populations). However, there was a correlation between the expression profiles of immune cells, leading in some instances to mixtures of PBMCs and HEK293T cells.

With few exceptions (Chromium), significantly fewer cells mapped to the mouse genome (half that of human cells, on average), leading to poorer clustering performance. However, the expected subpopulation composition of the colon was maintained overall. A small set of putative intestinal stem cells (Lgr5 and Smoc2 expression) were close (in transcriptional space) to rapidly proliferating transit amplifying (TA) cells (showing high ribosomal genes). Secretory cells (e.g. Muc2, Tff3, Agr2) resulted in a well-defined cluster. Enterocytes were more heterogeneous and ordered along their grade of lineage commitment. Notably, in some experiments two distinct clusters of enterocytes were identified, as well as a very small group of enterocyte progenitors. In addition to colon cells, fibroblasts and immune-cells were detected in all samples.

### Reference datasets

To compare the efficiency of scRNA-seq protocols in describing the structure of a mixed population, we produced a reference dataset with 30,807 human and 19,749 mouse cells. Cells were clustered and annotated as described above. Due to the high number of cells, major cell types were clustered and clearly identifiable using population marker genes (**Supplementary Fig.2a-b**). However, to improve cell-to-cell annotations, we combined clustering with additional analyses. To annotate human blood cells, we used *matchSCore2* (see Methods) using an annotated set of 2700 PBMCs^15^ as reference (**Supplementary Fig.2c-d**). We used cluster-specific markers of annotated populations as input to create a multinomial logistic model according to the *matchSCore2* algorithm. For each unknown cell, we assigned probability values for any possible cell identity, and the most likely identity was used for the classification (where this probability was >0.5; otherwise the cell was considered unclassified). Cell identities inferred by *matchSCore2* were highly consistent with clusters, with agreement ranging from 96% for CD4 T-cells to 100% for B-cells. Cell-by-cell prediction helped to identify smaller cell subsets, such as FCGR3A monocytes, dendritic cells and megakaryocytes. For all clusters, 17% of the cells remained unclassified (**Supplementary Fig.2c**). Half of these were previously annotated as HEK293T cells, which split into three different clusters because they varied in number of genes (**Supplementary Fig.2d**). Cells with fewer genes (cluster HEK293T cell2 and partially HEK293T cell3) were classified as CD4+ T-cells, although these did not show expression of any of the key blood markers. For the purposes of subsequent analysis, we removed the *unclear* cluster, representing 1% of the total number of cells, as well as the unclassified cells (except cells in HEK293T clusters). To further validate annotations, we assigned a score to each cell, corresponding to the overall expression of cell type signatures from the list of the top 100 computational markers (**Supplementary Fig.2d**). Transcriptional signatures revealed a set of cells from the HEK293T cell1 and HEK293T cell2 clusters showing high scores (>0.5, range 0-1)for multiple signatures. We considered these as potential doublets, and removed them. The remaining cells were then used to compute an unbiased set of cell-type specific markers.

In the case of the mouse reference sample, we used clustering to dissect the colon subpopulation structure (excluding immune cells and fibroblasts). The largest cluster was formed by immature enterocytes (**Supplementary Fig.3a-b**). Other clusters included similar proportions of mature enterocytes, secretory cells, transit-amplifying cells and other undifferentiated cells. To refine annotations of immature cells, we ordered cells by intermediate states and projected them along a trajectory (see Clustering). The trajectory analysis (**Supplementary Fig.3c-d**) revealed 9 different states, ranging from intestinal stem cells and transit-amplifying cells (expressing high levels of Lgr5, Smoc2, Top2a) to enterocytes (Slc26a3, Saa1). Based on the pseudo-ordering and expression levels of previously described markers, states were merged into four major groups (**Supplementary Fig.3d**). For annotation, we labeled these four groups as Intestinal Stem cells (ISC), Transit Amplifying cells (TA), Enterocyte progenitors (Epr), and Enterocyte (E). We combined this finer-grained annotation with the remaining cell types, and then computed population-specific gene markers for training the reference model.

#### MatchSCore2

To systematically compare cell types from the analysis of different methods, we used *matchSCore2*, a mathematical framework for classifying cell types based on reference data (https://github.com/elimereu/matchSCore2). The reference data consists of a matrix of gene expression counts in individual cells whose identity is known. The following preliminary steps were applied before training the model:

- *Normalization of the reference data*

Gene expression counts are log-normalized for each cell using the natural logarithm of 1 + counts per 10,000. Genes are then scaled and centered using the ScaleData function in the Seurat package.

- *Definition of signatures and their relative scores*

For each of the identity classes in the reference data, positive markers were computed using the Wilcoxon Rank sum test. The top 100 ranked markers in each class were used as the signature for that class. To each cell, we assigned a vector x=(x1,..,xn) of signature scores, where n is the number of classes in the reference sample. The *i-*th signature score is computed as follow:

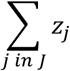

where J is the set of genes in signature *i*, and zj represents the z-score of gene *j*.

#### Statistical model

We proposed a supervised multinomial logistic regression model to explicitly infer cell identities. The model learns by training with the reference dataset, where n cell types and relatively ranked markers are defined. We assume that the distribution of signature scores is preserved, independent of which technology is used. The notion behind this model is that the random variable X=(X_1_,…,X_n_), where X*i* is the score in signature *i* across all cells, follows a multinomial distribution *M*(*s*= X_1_+..+ X_n_, π=(π_1_,…, π_n_)), where π*_i_* represents the proportion of the *i-th* cell type in the training set. Training and test sets were created by subsampling the reference into two datasets, maintaining the original proportions of cell types in both sets. The model was trained by using the *multinom* function from the *nnet* R package (*decay*=1e-04, *maxit*=500). To improve the convergence of the model function, X*_i_* variables were scaled to the interval [0,1].

#### Cell Classification

For each cell, model predictions consisted of a set of probability values per identity class, and the highest probability was used to annotate the cell if it was >0.5; otherwise the cell resulted unclassified.

#### Model accuracy

To evaluate the fitted model using our reference datasets, we assessed the prediction accuracy in the test set, which was around 0.9 for human and 0.85 for mouse reference. We further assessed *matchSCore2* classifications in datasets from other sequencing methods by looking at the agreement between clusters and classification. Notably, the resulting average agreement was of 80% (range: from 58% in gmcSCRB-seq to 92% in Quartz-Seq2), while the rate of unclassified cells was less than 2%.

### Downsampling

To decide on a common downsampling threshold for sequencing depth per cell, we inspected the distribution of the total number of reads per cell for each technique, and chose the lowest first quartile (fixed to 20,000 reads/cell). We then performed stepwise downsampling (25%, 50% and 75%) using the zUMIs downsampling function. We omitted cells that did not achieve the required minimum depth.

### Estimation of dropout probabilities

We investigated the impact of dropout events in HEK293T, monocytes and B-cells extracted for each technique on downsampled data (20,000 reads/cell). For datasets with >50 cells from the selected populations, we randomly sampled 50 cells to eliminate the effect of differing cell number. The dropout probability was computed using SCDE R package^42^. SCDE models the measurements of each cell as a mixture of a negative binomial process to account for the correlation between amplification and detection of a transcript and its abundance, and a Poisson process to account for the background signal. We then used estimated individual error models for each cell as a function of expression magnitude to compute dropout probabilities using SCDE’s scde.failure.probability function. Next, we calculated the average estimated dropout probability for each cell type and technique. To integrate dropout measures into the final benchmarking score, we calculated the Area Under the Curve (AUC) of the expression prior and failure probabilities (Figure 2f). We expected that protocols that result in fewer dropouts would have lower AUC.

### Cumulative number of genes

The cumulative number of detected genes in downsampled data was calculated separately for each cell type. For cell types with >50 cells annotated, we randomly selected 50 cells and calculated the average number of detected genes per cell after 100 permutations over n sampled cells, where n is an increasing sequence of integers from 1 to 50.

### Silhouette scores

To measure the strength of the clusters, we calculated the Average Silhouette Width (ASW)^18^. The downsampled data (20,000 reads/cell) were clustered by Seurat^40^, using graph based clustering with the first 8 principle components and resolution of 0.6. We then computed an average Silhouette width for the clusters using an Euclidean distance matrix (based on principle components 1 to 8). We report the average Silhouette width for each technique separately.

### Dataset merging

Dataset integration across studies is one of the most challenging analyses. It is important to assess which scRNA-seq methods integrate best, while conserving biological variability. To integrate datasets, we used the R package *scMerge*^19^, which uses a set of genes with stable expression levels across different cell types. Also, creating pseudo-replicates across datasets allows to estimate and correct for undesired sources of variability. To avoid differences due to sequencing depth, we combined downsampled count matrixes using the *sce_cbind* function, which includes the union of genes from different batches. After computing the set of highly variable genes using log-normalized gene measurements, we then apply the *scmerge* function to align data from different experiments. Following integration, cells were clustered using normalized gene expression levels and HVG computed using scMerge. We used UMAP plots color-coded by clusters and cell types to visualize and annotate clusters with the greatest agreement between cell types and clustering.

### Clustering accuracy

To determine the clusterability of methods to identify cell types, we measured the probability of cells to be clustered with cells of the same type. Let *C*_k_, *k* ∈ {1, …, *N*} the cluster of cells corresponding to a unique cell type (based on the highest agreement between clusters and cell types), and *T*_*j*_, *j* ∈ {1, …, *S*} the set of different cell types, where C⊆T. For each cell type Tj, we compute the proportion pjk of Tj cells that cluster in their correct cluster Ck. We define the cell-type separation accuracy as the average of these proportions.

### Mixability

To account for the level of mixing of each technology, we used kBet^20^ to quantify batch effects by measuring the rejection rate of a Pearson’s χ2 test for random neighborhoods. To make a fair comparison, kBet was applied to the common cell types separately by subsampling batches to the minimum number of cells in each cell type. Due to the reduced number of cells, the option heuristic was set to ‘False’, and the testSize was increased to ensure a minimum number of cells. Mixability was calculated by averaging cell type specific rejection rates.

### Benchmarking score

To create an overall benchmarking score with which to compare technologies, we considered six key metrics: gene detection, overall level of expression in transcriptional signatures, cluster accuracy, classification probability, cluster accuracy after integration, and mixability. Each metric was scaled to the interval [0,1], then in order to equalize the weight of each metric score, the harmonic mean across these metrics was calculated to obtain the final Benchmarking scores. Gene detection, overall expression in cell type signatures, and classification probabilities were computed separately for B-cells, HEK293T cells and monocytes, and then aggregated by the arithmetic mean across cell types.

