## Supplementary Material for "Benchmarking Single-Cell RNA Sequencing Protocols for Cell Atlas Projects"

**Supplementary Figure legends 1-11.**

**Supplementary Figures 1-11.**

**Supplementary Table 1.**

**Supplementary Fig. 1. Gene expression levels of selected marker genes.**
UMAP visualization of normalized expression levels for selected marker genes of the most common PBMC (**a**) and colon (**b**) populations. Maps are shown for CD4+ T-cell markers IL7R and CD4 (expressed also in monocytes), the CD8+ T-cell marker CD8A, the B-cell marker CD79A, NK cell markers GNLY and NKG7, and monocyte-specific markers LYZ, CD14 and FCGR3A. In (**b**) maps are shown for markers of Intestinal Stem cell and proliferation (Smoc2, Miki67 and Top2a), secretory markers (Muc2, Agr2 and Tff3), enteroendocrine cell markers (Chga and Chgb), and enterocyte markers (Slc26a3, Car1 and Fabp2).

**Supplementary Fig. 2. Identifying PBMC cell types using unsupervised clustering and classification. a.** UMAP visualization of 38,195 human PBMC and HEK293T human cells coloured according to their assignment to clusters. Cluster labels are defined by examining the expression levels of known markers. **b.** Heatmap indicating the relative expression and gene detection rates for most common PBMC marker genes. **c.** UMAP vizualization of PBMC and HEK293T cells colour coded by cell classification inferred by matchSCore2. 17% of cells were unclassified and were removed from the analysis. **d.** UMAP visualization of cells showing the number of genes per cell, and scores for transcriptional signatures obtained by computing cell-type-specific markers (*lightgray*: low-score, *blue*: high score).

**Supplementary Fig. 3. Identifying colon cell types by unsupervised clustering and trajectory analysis. a.** UMAP visualization of 17,558 mouse colon cells. Cells are coloured by their assignment to clusters. Annotations are defined by examining the expression of known markers and differentially expressed genes (DEG). **b.** Heatmap of top DEG per cluster. Key markers of common colon cell populations are shown. **c.** Trajectory and pseudotime analysis of 8716 immature enterocytes (IE) showing the transition from intestinal stem cells (ISC) to enterocytes. Trajectories with the relative expression of known markers are shown (yellow: low, gray: mid, blue: high). **d.** (Top) Ordered cells are grouped into four different states according to their differentiation stage: intestinal stem cell (ISC), transit amplifying (TA), enterocyte progenitor (Epr), Enterocytes (E). (Bottom) UMAP visualization of IE cells coloured according to the four resulting states.

**Supplementary Fig. 4. Clustering analysis of 13 sc/snRNA-seq methods.** T-SNE visualizations of unsupervised clustering in human samples from 13 different methods. Each dataset is analyzed separately by taking advange of its original sequencing depth. Cells are coloured by cell type inferred by *matchSCore2*. Cells that did not reach a probability score of 0.5 for any cell type were considered unclassified.

**Supplementary Fig. 5. Clustering analysis of 11 sc/snRNA-seq methods.** T-SNE visualizations of the unsupervised clustering in mouse samples from 11 different methods. Each dataset is analyzed separately by taking advange of its original depth. Cells are coloured according to cell type inferred by *matchSCore2*. Cells that did not reach a probability score of 0.5 for any cell types were considered unclassified.

**Supplementary Fig. 6. Comparison of 13 scRNA sequencing methods in mouse data.**
**a.** Boxplots comparing the number of detected genes across protocols on downsampled data (20K), in mouse secretory and transit-apmlifying cells. Cell identities were defined by cell projection onto the reference**. b.** Number of genes detected at step-wise downsampled sequencing depths. Points represent the average number of genes detected for all cells of the corresponding cell type at the corresponding sequencing depth. **c.** Boxplots comparing the number of detected genes from countification of reads mapping to only Exonic regions, across protocols on downsampled data (20K), in Human HEK293T cells, monocytes and B-cells.

**Supplementary Fig. 7. T-SNE representation of human cell types using highly variable genes.**
**a,b.** T-SNE representation (calculated on first 8 principle components) on downsampled data (20K) using highly variable genes across protocols, separated by HEK293T cells, monocytes and B-cells and color coded by protocols (**a**) or the number of detected genes per cell (**b**).

**Supplementary Fig. 8. PCA representation of Human cell types using cell type markers**
**a,b.** PCA analysis on downsampled data (20K) for HEK293T cells, monocytes and B-cells separately using the corresponding cell type's reference markers and color coded by protocols (**a**) or number of detected genes per cell (**b**).

**Supplementary Fig. 9. Gene expression correlations across 13 sc/snRNA-seq methods.** Pearson correlation plots between protocols using gene expression of cell-type-specific signatures for HEK293T cells (**a**), monocytes (**b**) and B-cells (**c**). Cells are ordered by agglomerative hierarchical clustering.

**Supplementary Fig. 10. Analysis of integrated methods. a,b.** UMAP visualization of clusters after the integration of technologies for human (**a**) and mouse (**b**) datasets. Cluster annotations are assigned on the basis of the most frequent cell type. **c,d.** Barplots showing normalized and method-corrected (integrated) expression scores in cell type specific signatures for CD4+ and CD8+ T-cells (**c**) and enterocytes 1, enterocytes 2 and intestinal stem cells (**d**). Bars are coloured by method. **e.** Evaluation of method integrability. Protocols are compared in their ability to group cell types into clusters (after integration) and to mix with other technologies within same clusters. Point sizes are indicating the level of downsampled sequencing depths. Dotted lines connect points from the same technology, highlighting the drop of integratability at lower depth. Points are coloured by sequencing method.

**Supplementary Fig. 11. Comparison of mappability scores across technologies.** Boxplots displaying minimum, 1st, 2nd, 3rd quantiles and maximum probabilities values (scores) obtained by *matchSCore2* in classifying most common cell types in human (**a,b**) and mouse (**c**) samples. B-cells, HEK293T cells and CD14+ monocytes are shown with data downsampled to 20K (**a**) and 10K (**b**) sequencing reads.

Supplementary Figure 1

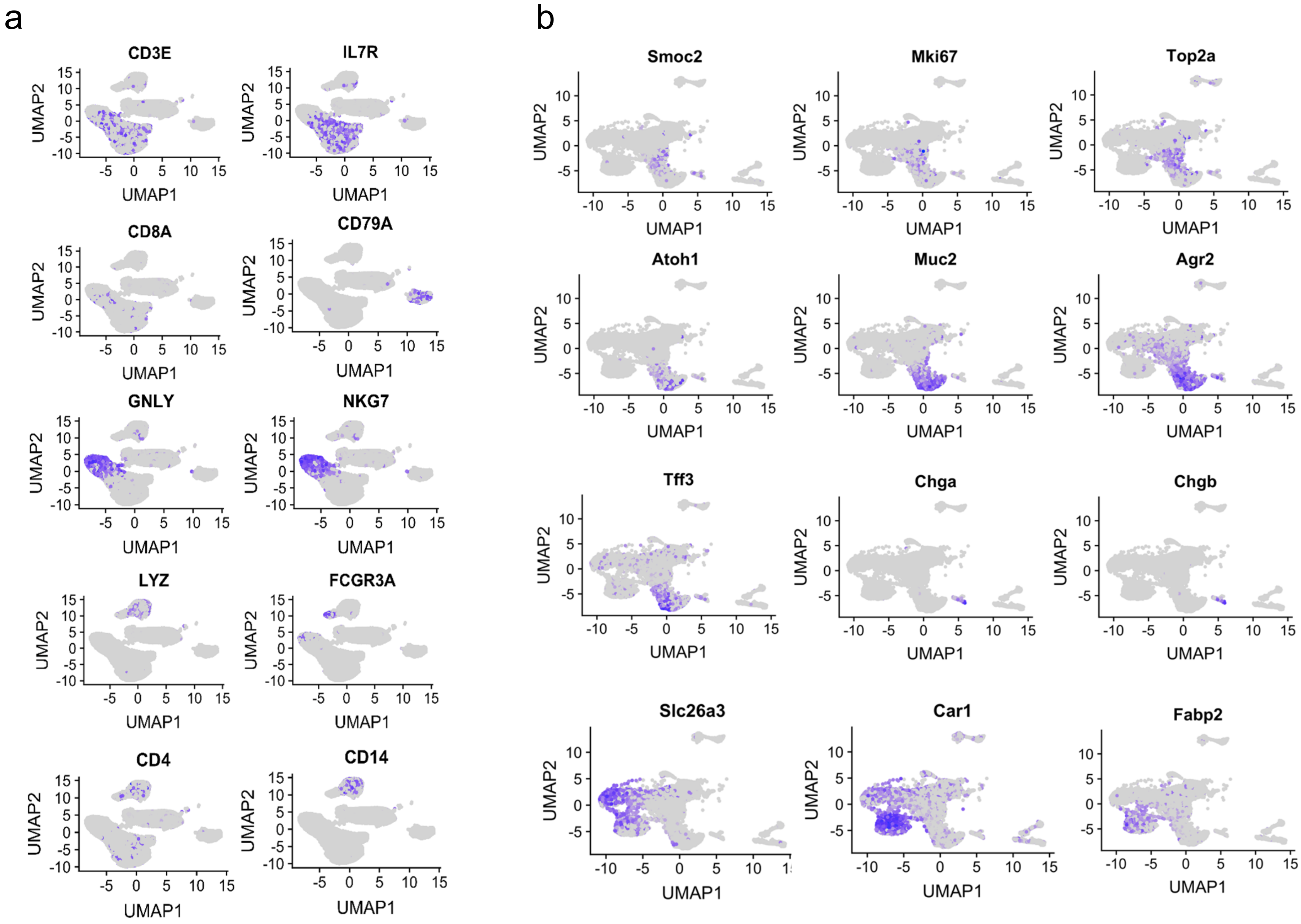

Supplementary Figure 2

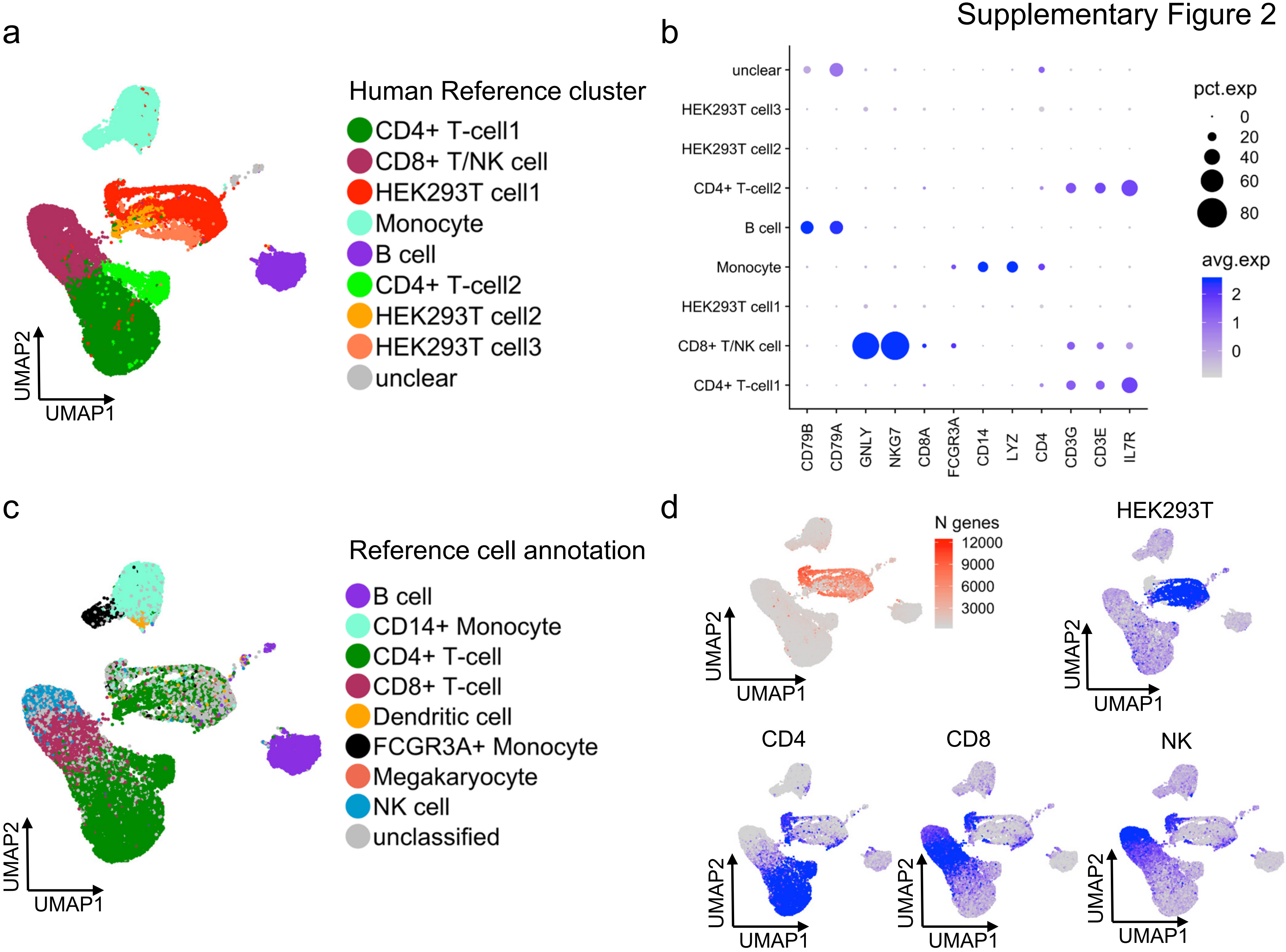

Supplementary Figure 3

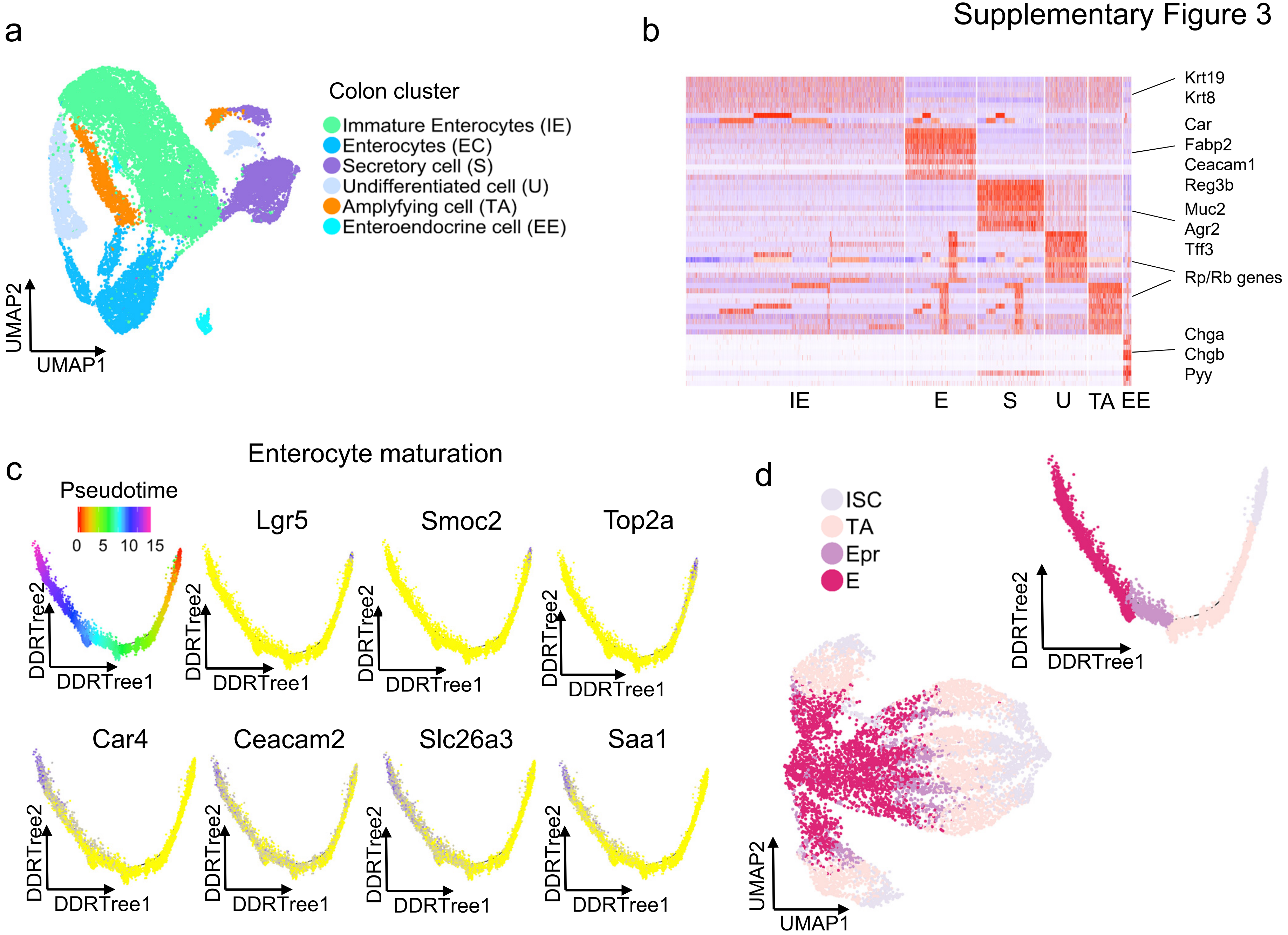

Supplementary Figure 4

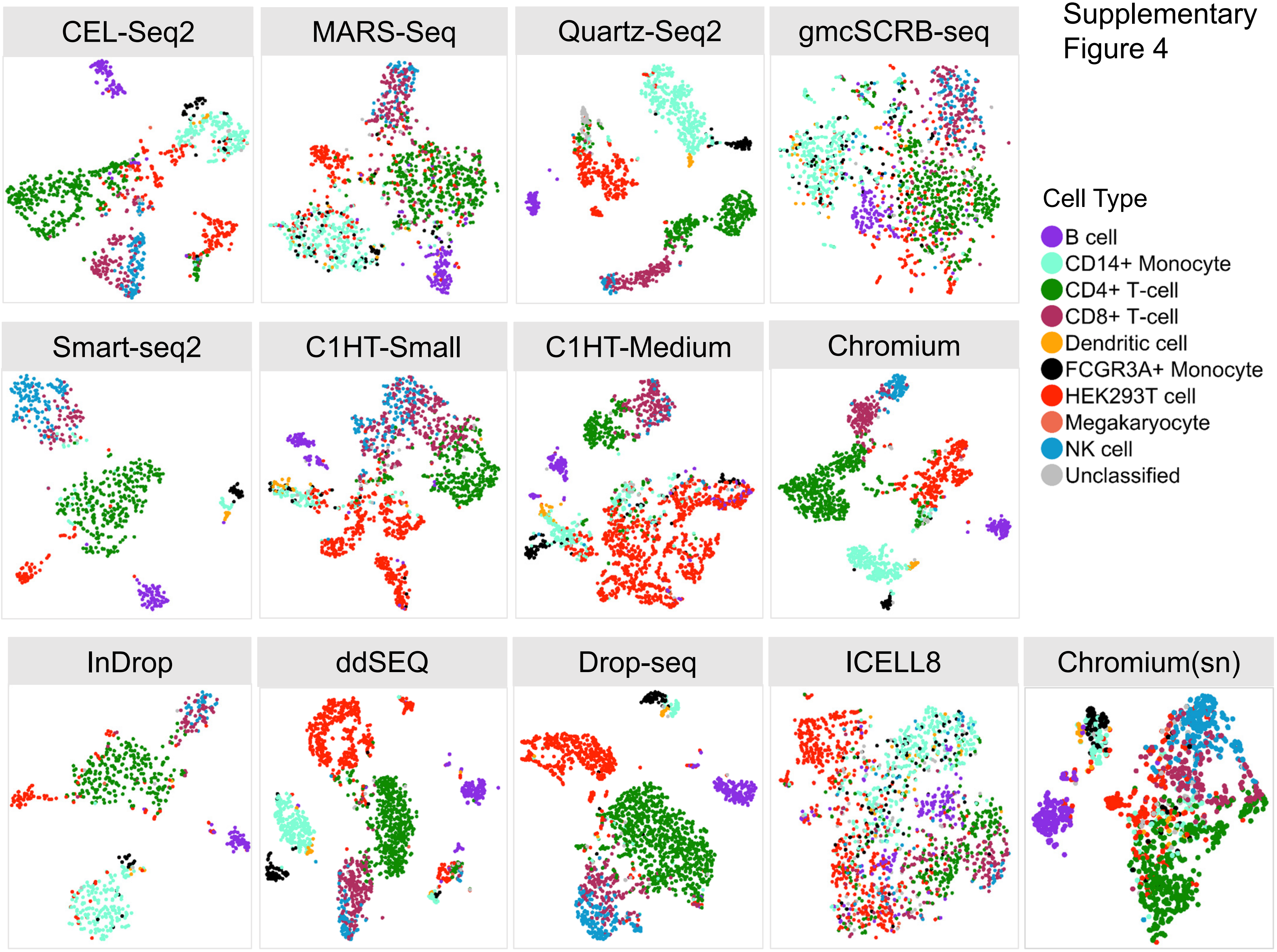

Supplementary Figure 5

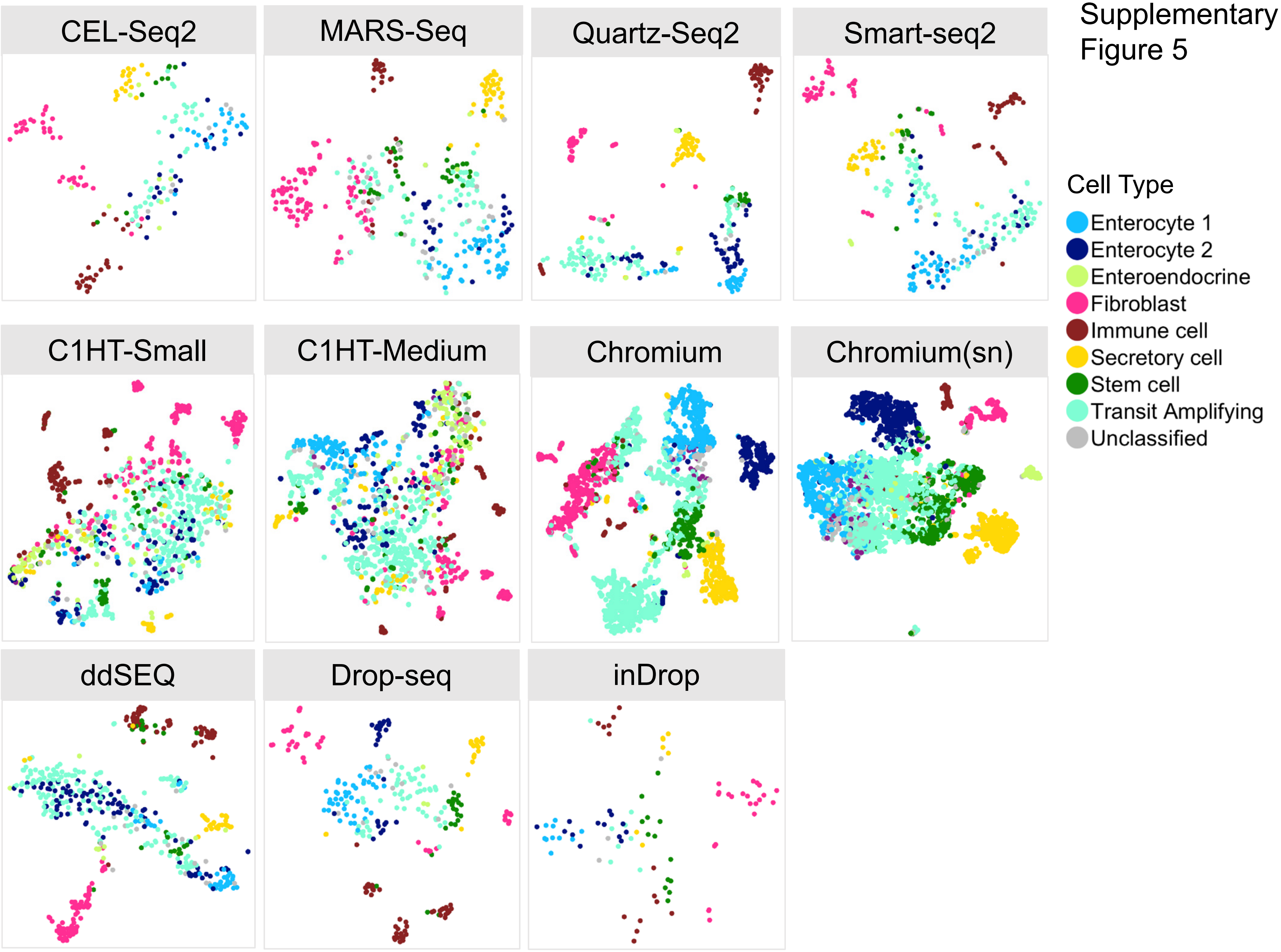

Supplementary Figure 6

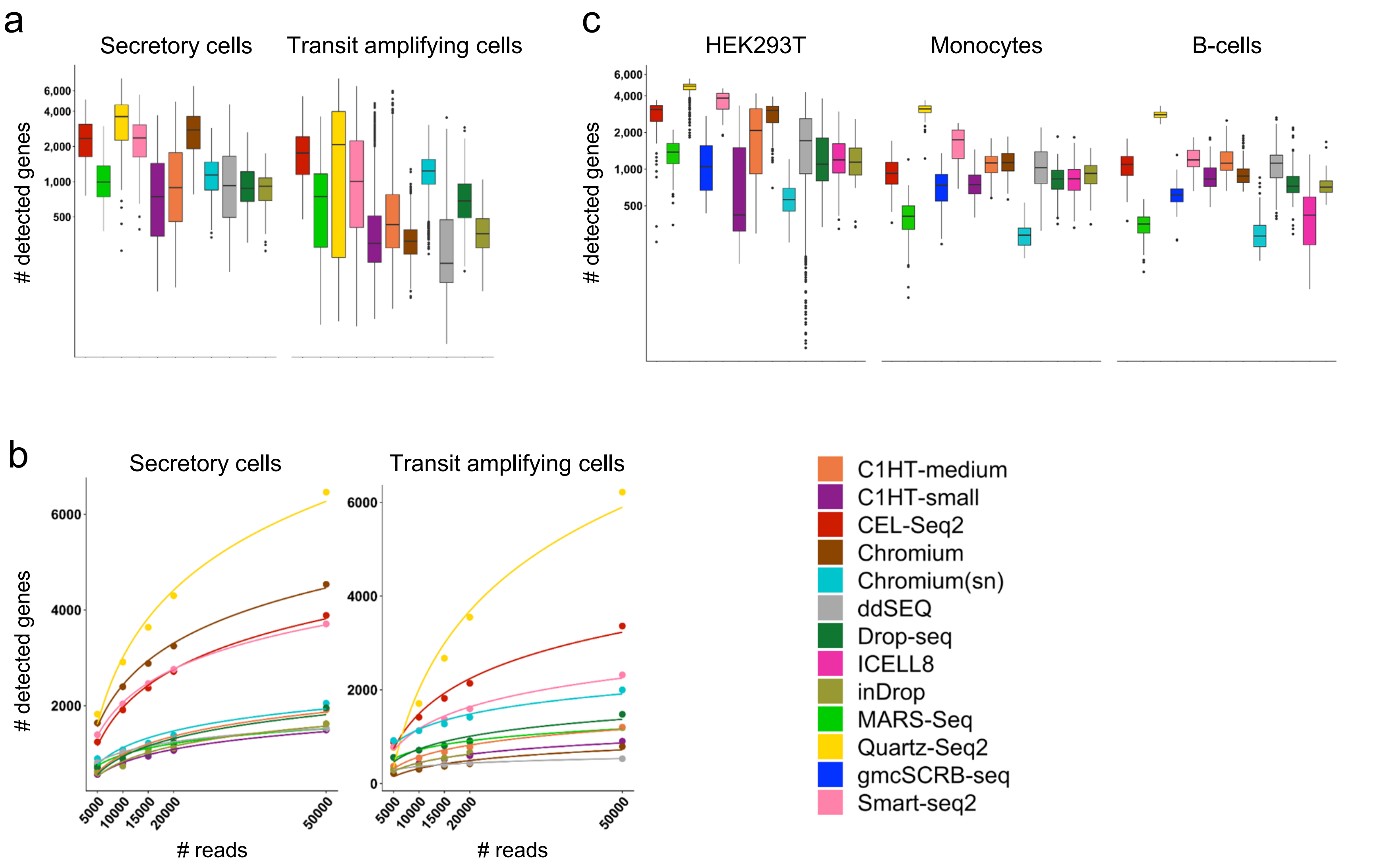

Supplementary Figure 7

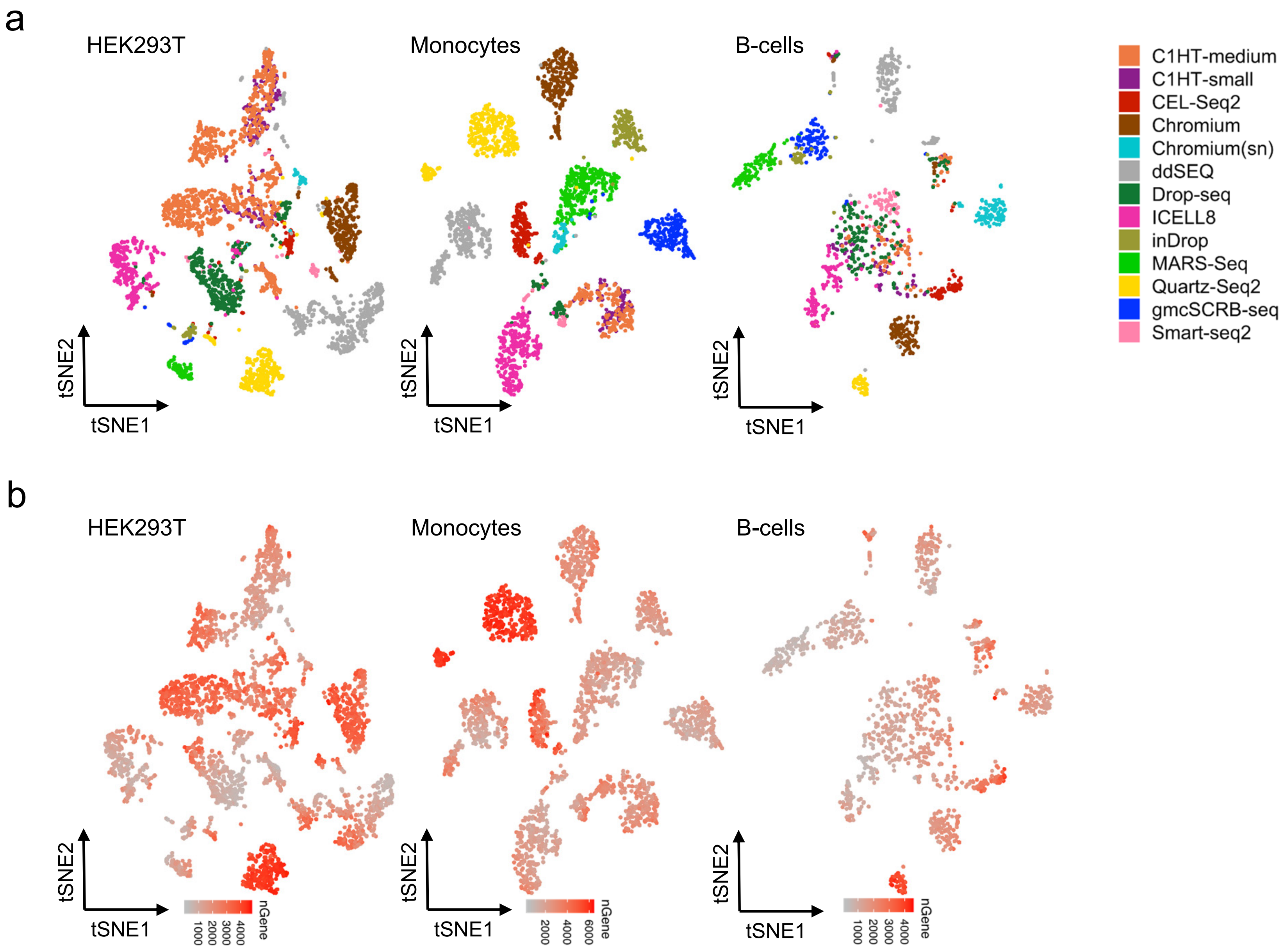

Supplementary Figure 8

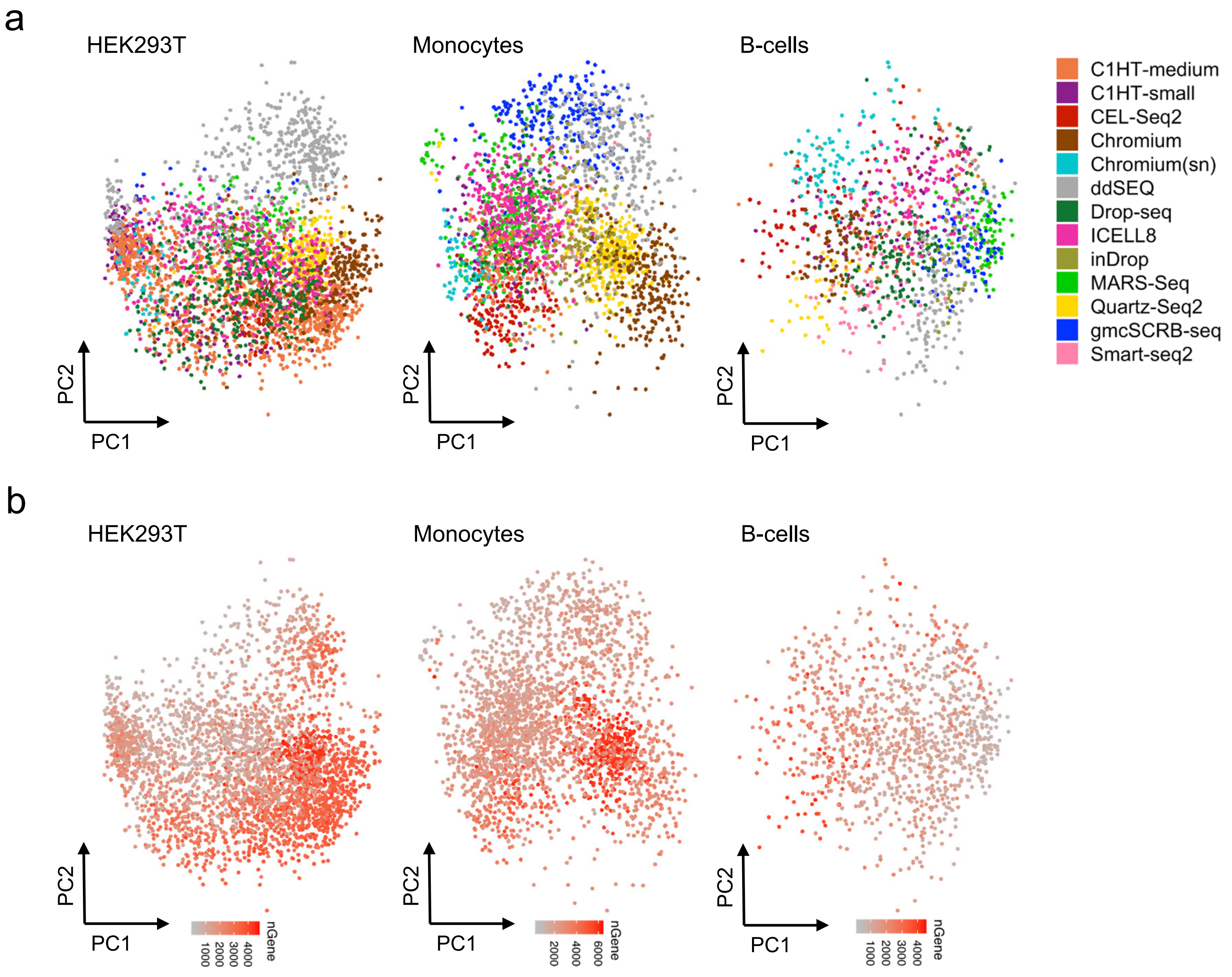

Supplementary Figure 9

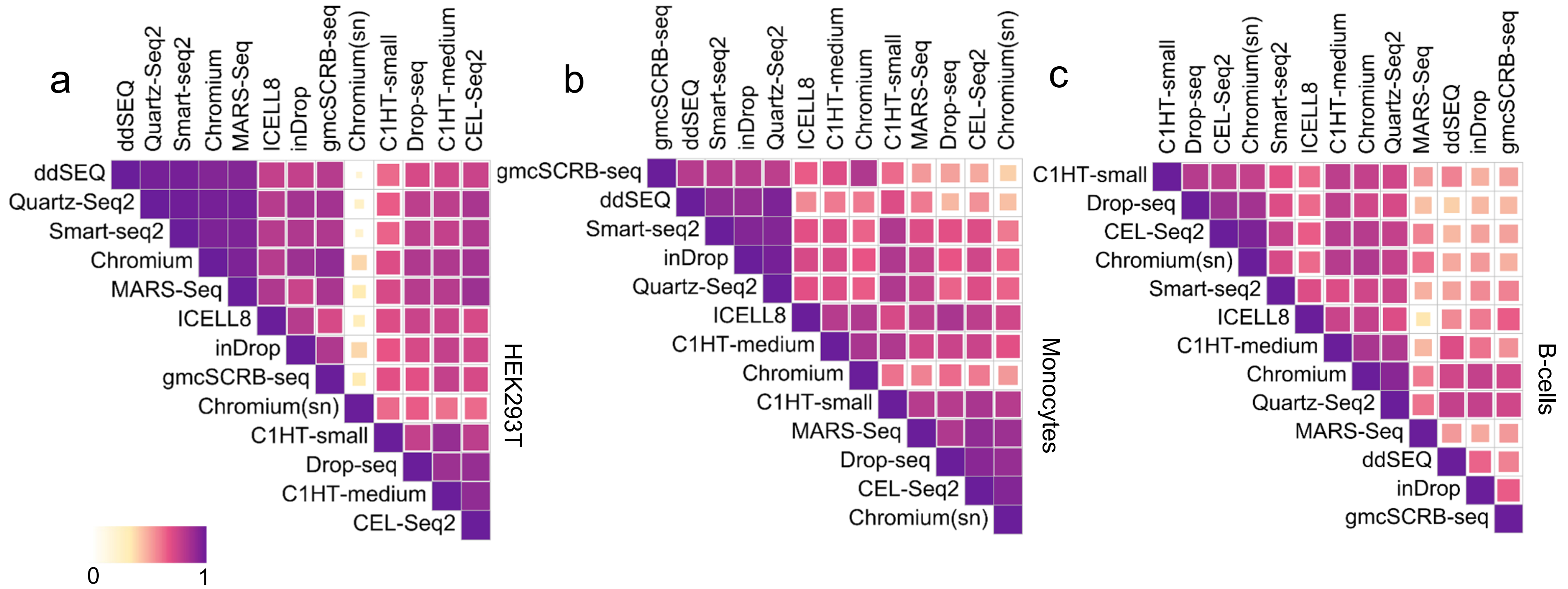

Supplementary Figure 10

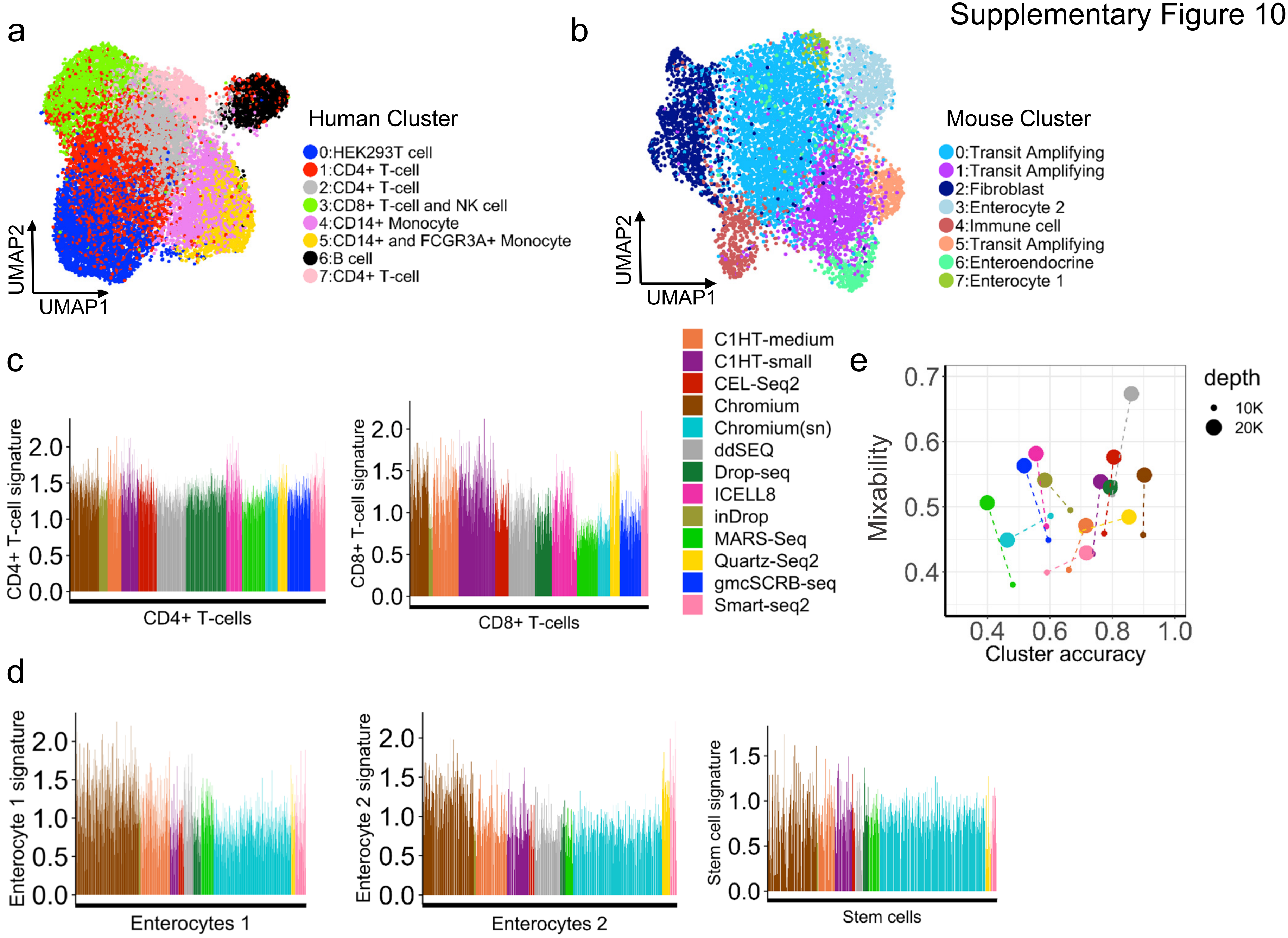

Supplementary Figure 11

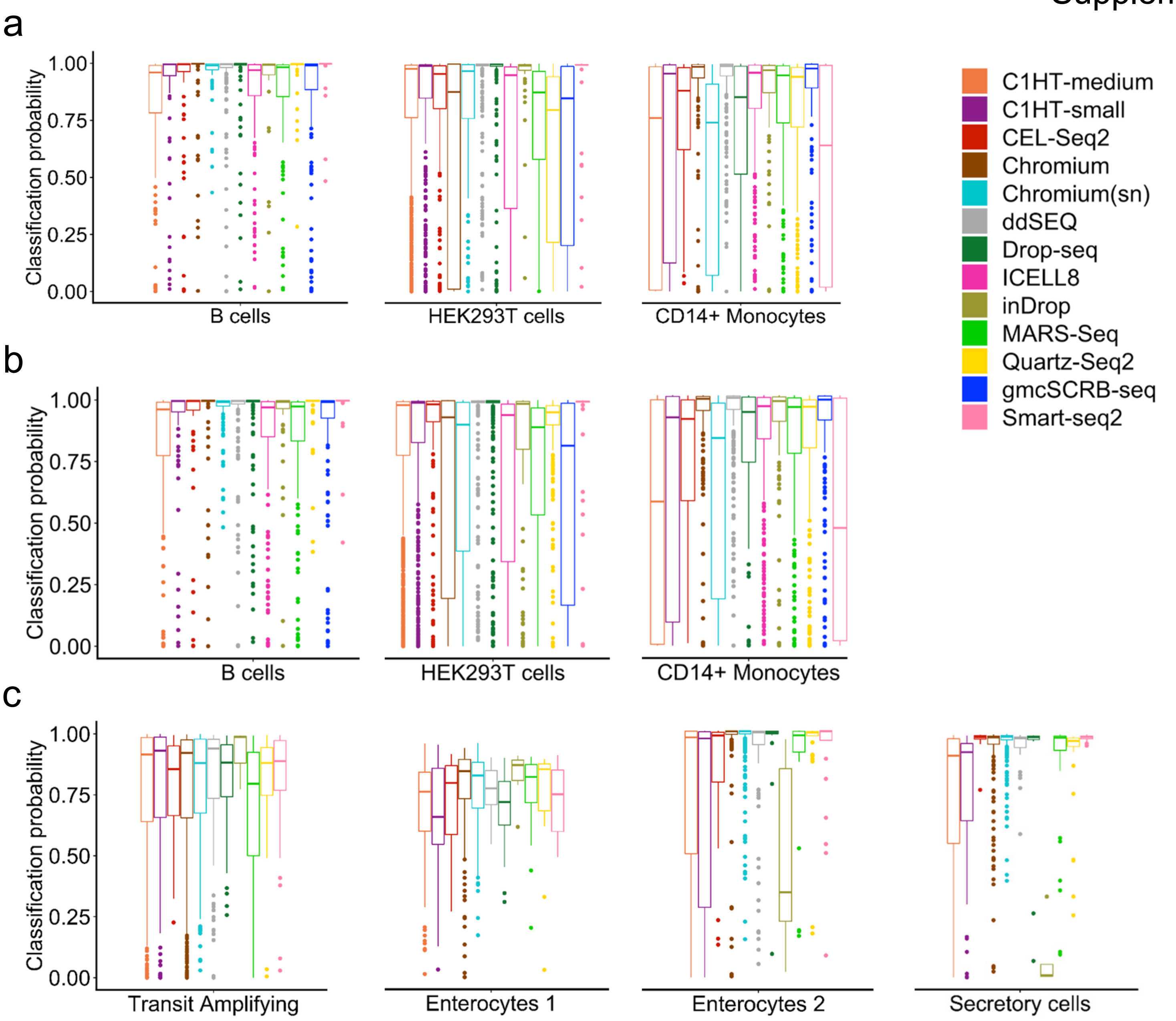

**Table S1:** Single-cell RNA sequencing methods and protocol specific features.

| Method | Provider | Capture format | Cell loading | Single-cell indexing | Molecule identifier | cDNA amplification | Transcript coverage | Ref. |
|---|---|---|---|---|---|---|---|---|
| C1HT | Fluidigm | IFC | Trapping | OligoT primer | UMI | PCR | 3´-end | [1] |
| CEL-Seq2 | | Plate | FACS | OligoT primer | UMI | IVT | 3´-end | [2,3] |
| Chromium | 10x Genomics | Droplets | Poisson | OligoT beads | UMI | PCR | 3´-end | [4] |
| ddSEQ | Bio-Rad | Droplets | Double Poisson | OligoT beads | UMI | PCR | 3´-end | [5] |
| Drop-seq | Dolomite | Droplets | Double Poisson | OligoT beads | UMI | PCR | 3´-end | [6] |
| ICELL8 | Takara Bio | Nanowells | Poisson | OligoT probes | UMI | PCR | 3´-end | [7] |
| inDrops | 1CellBio | Droplets | Poisson | OligoT beads | UMI | IVT | 3´-end | [8] |
| gmcSCRB-seq | | Plate | FACS | OligoT primer | UMI | PCR | 3´-end | [9] |
| MARS-Seq | | Plate | FACS | OligoT primer | UMI | IVT | 3´-end | [10] |
| Quartz-Seq2 | | Plate | FACS | OligoT primer | UMI | PCR | 3´-end | [11] |
| Smart-seq2 | | Plate | FACS | Tagmentation | N/A | PCR | Full-length | [12] |

### Table S1 References
